# Chemical Probing of Diacylglycerol Dynamics at Lipid Droplets

**DOI:** 10.64898/2026.02.27.708637

**Authors:** Christopher Gomez, Nicholas McInchak, Sarah Louis, Evan Heck, Haylee Mesa, Jonathan Meade, Qi Zhang, Maciej J. Stawikowski

**Affiliations:** Department of Chemistry and Biochemistry, Charles E. Schmidt College of Science, Florida Atlantic University, 777 Glades Rd., Boca Raton, FL 33431, USA; Department of Chemistry and Biochemistry, The Stiles-Nicholson Brain Institute, Florida Atlantic University, 5353 Parkside Dr., Jupiter, FL 33458, USA

## Abstract

Diacylglycerols (DAGs) are central intermediates in lipid metabolism and signaling, yet their trafficking and persistence within lipid droplets (LDs) remain incompletely understood due to the lack of chemically stable, DAG-mimetic imaging tools. Here, we report the development of a family of solvatochromic fluorescent lipid analogs, termed DONDI, designed to probe DAG-associated dynamics at LDs. These probes are based on a 1,8-naphthalimide scaffold conjugated to modified aminoglycerol backbones bearing oleoyl chains to mimic native glycerolipids. Biophysical characterization and atomistic molecular dynamics simulations revealed probe-specific membrane insertion and hydrogen-bonding behaviors consistent with distinct lipid-mimetic properties. Live-cell imaging in NIH 3T3 fibroblasts demonstrated that DONDI probes were efficiently internalized and selectively accumulated within lipid droplets. Structure–function analysis identified DONDI-5 as the closest mimic of 1,2-diacylglycerol, displaying rapid uptake, strong LD enrichment, and prolonged intracellular retention without detectable relocalization to other cellular membranes. These properties enabled sustained visualization of LD-associated DAG pools over extended time scales. Collectively, this work establishes DONDI-5 as a chemically stable DAG-mimetic probe and provides direct experimental support that DAGs can be transported to and transiently stored within lipid droplets without prior conversion to triacylglycerols.

## INTRODUCTION

Neutral glycerolipids, including diacylglycerols (DAGs) and triacylglycerols (TAGs), occupy central positions in cellular lipid metabolism.^1^ Their shared glycerol backbone provides a versatile scaffold that supports the synthesis and interconversion of multiple lipid classes, including serving as intermediates in glycerophospholipid biosynthesis. The reversible conversion between TAGs and DAGs is tightly regulated by lipases and acyltransferases, enabling cells to dynamically balance lipid storage and utilization. Importantly, TAG hydrolysis generates structurally distinct DAG species. Because TAGs are prochiral, lipase-mediated cleavage can yield achiral (racemic) 1,3-DAG or chiral *sn-*1,2- and *sn-*2,3-DAG isomers. These regio- and stereochemical variants are not functionally equivalent: beyond their metabolic roles, DAGs act as key secondary messengers that engage distinct downstream signaling pathways. Consequently, the spatial and structural handling of DAG species represents a critical node at the intersection of lipid metabolism and cell signaling. Intracellular lipid droplets (LDs), which reside in the cytosol and maintain extensive contacts with the endoplasmic reticulum (ER), function as major repositories for neutral lipids, including TAGs, DAGs, and cholesteryl esters (CEs).^2^ LDs actively regulate lipid flux to and from other organelles to support energy homeostasis, membrane biogenesis, and the generation of signaling lipids derived from fatty acid precursors. In addition, LDs serve a protective role by sequestering excess or potentially lipotoxic lipid species.^3^ Despite their central metabolic importance, the molecular features governing DAG distribution, persistence, and remodeling within LDs remain incompletely understood. Given the pivotal roles of DAGs in both signaling and neutral lipid metabolism^4,5^, we sought to develop fluorescent DAG analogs that report DAG-like behavior within lipid droplets. Such probes would provide a means to interrogate LD-associated lipid dynamics, including biogenesis, turnover, and lipid exchange at membrane contact sites, while preserving key structural features relevant to endogenous DAG function. Notably, a key challenge in developing functional DAG probes is preserving the defining free hydroxyl group at the *sn-*3 position, a structural feature critical for DAG recognition and behavior that is frequently compromised by fluorophore conjugation strategies. Accordingly, we sought to develop fluorescent DAG analogs that preserve key structural features of DAGs while incorporating a compact, environment-sensitive fluorophore compatible with lipid-rich cellular compartments.

## RESULTS AND DISCUSSION

### Design and synthesis of fluorescent probes

We designed and synthesized four novel diacylglycerol probes featuring an amino glycerol scaffold conjugated to a 1,8-naphthalimide fluorophore (Figure 1). The core fluorophore moiety is covalently linked to the lipid backbone *via* a carbon-nitrogen (C-N) bond, requiring the use of an aminoglycerol as a precursor. Among the commercially available aminoglycerol derivatives, we identified two suitable scaffolds for constructing fluorescent DAG probes: chiral (*2S*)-3-amino-1,2-propanediol and prochiral 2-amino-1,3-propanediol. To account for the stereochemical requirements of the *sn*-glycerol backbone, we selected (*2S*)-3-amino glycerol for the synthesis of DONDI-1, DONDI-3, and DONDI-4, and used the achiral 2-amino glycerol for DONDI-5.

**Figure 1.**
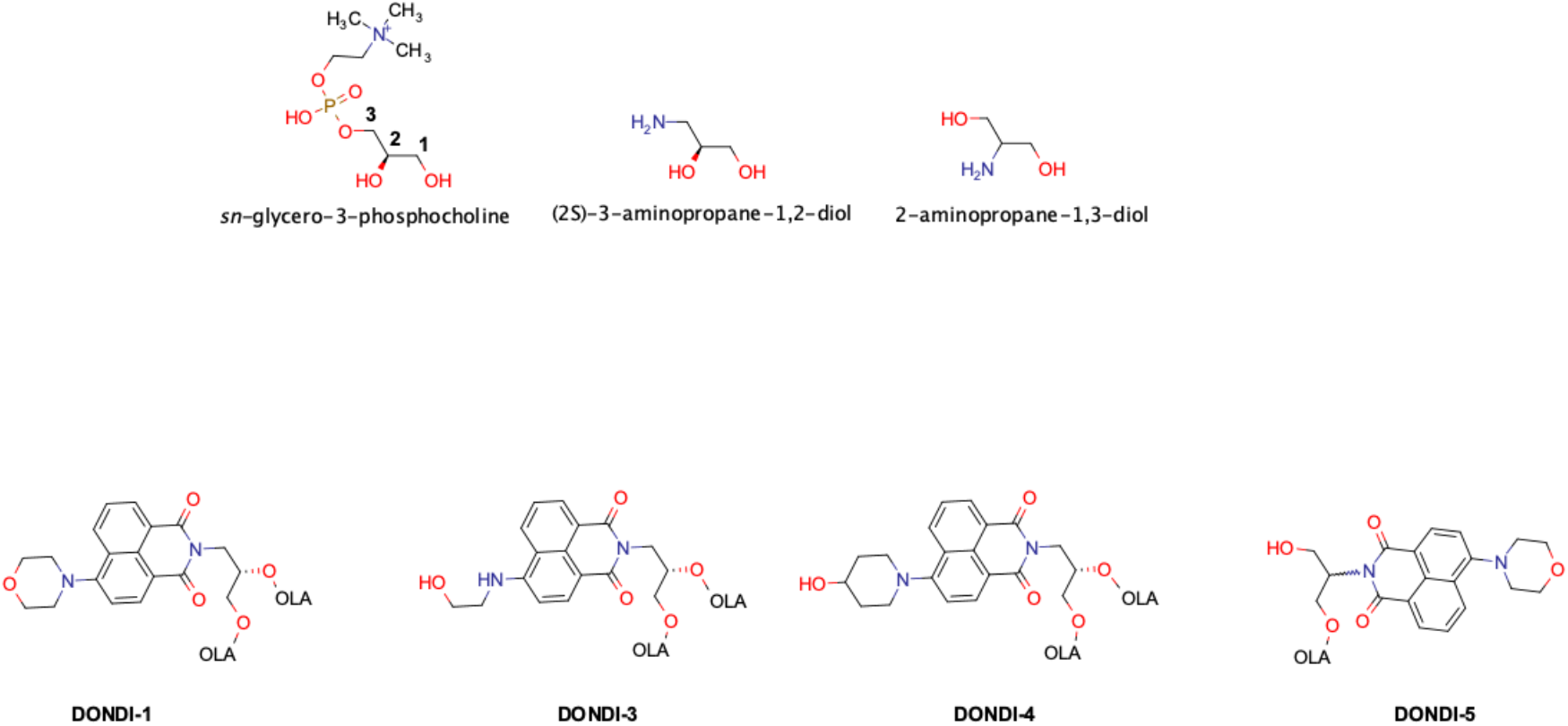
DONDI analogs investigated in this study. Top row: Comparison of aminoglycerol scaffolds used for probe design. Bottom row: Chemical structures of the corresponding DONDI analogs (DONDI-1, DONDI-3, DONDI-4, and DONDI-5). OLA - oleic acid

Native DAG and TAG species typically exhibit regioselective acylation, with saturated fatty acids preferentially occupying the *sn-*1 position and unsaturated chains at the *sn-*2 position. For proof-of-concept studies and to simplify structure–function comparisons, oleic acid (18:1) was used for acylation at all available hydroxyl groups. Initially, when stearic acid (18:0) was used, the resulting compounds exhibited poor solubility in common organic solvents, limiting their suitability for live-cell imaging. In contrast, oleic acid-containing probes showed dramatically improved solubility in dimethyl sulfoxide by approximately three orders of magnitude, enabling biological studies at relevant concentrations.

Previous work from our laboratory demonstrated that substitution at the C-4 position of the naphthalimide scaffold strongly influences both the fluorophore’s photophysics and its membrane interactions.^6^ Guided by this insight, we restricted head-group modifications to naturally occurring polar motifs - morpholine, ethanolamine, and 4-hydroxymorpholine - to preserve the biomimetic character of the probes. Notably, a defining structural feature of native DAGs is the presence of a free hydroxyl group at the *sn-*2 or *sn-*3 position, which is critical for interactions with DAG-responsive proteins. Although direct fluorophore conjugation at the *sn-*3 position necessarily masks this defining hydroxyl group, we reasoned that systematic variation in fluorophore placement and head-group polarity could provide insight into how such structural perturbations influence DAG-versus TAG-like behavior. In the structures of DONDI-1, DONDI-3, and DONDI-4, the fluorescent 1,8-naphthalimide moiety is covalently attached to the *sn*-3 carbon of the glycerol backbone (Figure 1). In native 1,2-diacylglycerols (DAGs) this position is occupied by a free hydroxyl group, thereby potentially altering the probes’ functional resemblance to endogenous DAGs. Interestingly, this substitution could mimic the architecture of triacylglycerols (TAGs), which have all three hydroxyl groups of glycerol esterified, suggesting that these probes may behave more like TAG analogs in biological systems rather than true DAG surrogates. To investigate this hypothesis and to more closely mimic the native structure of DAG - specifically, the presence of a free hydroxyl group at the *sn*-3 position - we designed an alternative analog, DONDI-5. In this analog, the naphthalimide fluorophore is instead conjugated to the *sn*-2 carbon of glycerol, leaving the *sn*-3 position unmodified and bearing a free hydroxyl group. By comparing the biophysical properties and biological behavior of DONDI analogs, we aimed to design functional fluorescent diacylglycerol analogs.

The synthesis of DONDI analogs was accomplished in a three-step reaction sequence (see Supporting Information). In the first step, the appropriate aminoglycerol scaffold was reacted with 4-bromo-1,8-naphthalic anhydride to form the corresponding naphthalimide intermediate (Scheme S1, compounds A1/B1). This reaction proceeds *via* nucleophilic attack of the amine on the anhydride, yielding the core fluorophore conjugated to the glycerol backbone. In the second step, the bromine atom at the C-4 position of the naphthalimide ring was displaced *via* nucleophilic aromatic substitution with the desired primary or secondary amine, yielding the functionalized naphthalimide-glycerol conjugates. These intermediates were isolated in moderate yields; however, their poor solubility in most organic solvents presented a challenge, often limiting reaction efficiency and purification. In the final step, the free hydroxyl groups were esterified with oleic acid (OLA), except for DONDI-5, which was esterified with only a single hydroxyl group. The synthetic details are provided in the supporting information section.

### Spectroscopic and biophysical characterization

The 1,8-naphthalimide fluorophore is well known for its pronounced solvatochromism, making it particularly suitable for environmentally sensitive fluorescent probes.^7,8^ This property is advantageous for lipid-associated reporters, as solvent-dependent spectral shifts can reflect changes in local polarity and microenvironment encountered upon membrane insertion or aggregation. To characterize the solvatochromic response of the DONDI analogs, UV–Vis absorption and fluorescence emission spectra were recorded in chloroform and dimethyl sulfoxide (DMSO), two solvents with substantially different polarities. As shown in Figure S1, all DONDI probes exhibited similar absorption profiles in chloroform, with characteristic absorption maxima near 400 nm. DONDI-3 displayed a modest red shift relative to the other analogs, consistent with subtle differences in head-group substitution. In DMSO, all probes showed a pronounced bathochromic shift compared to chloroform, reflecting the increased solvent polarity and consistent with prior reports on related naphthalimide-based fluorophores, including the cholesterol-derived CND probe series.^9,10^ Fluorescence emission spectra revealed large Stokes shifts (110–130 nm), a hallmark of naphthalimide fluorophores that minimizes self-quenching and enhances signal-to-noise ratios for fluorescence imaging. All DONDI analogs exhibited strong emission in the green spectral region, with maxima between 520 and 550 nm. When DONDI probes were introduced into aqueous phosphate-buffered saline (PBS) from DMSO stock solutions, substantial fluorescence was observed even in the absence of model membranes. This behavior may suggest a spontaneous self-association into fluorescent aggregates or micellar assemblies in aqueous (non-membrane) environments. Unlike previously reported cholesterol-based neutral CND probes (e.g., CND5, CND6, and CND9), these assemblies did not form visible precipitates, potentially reflecting differences in hydrophobic anchoring and molecular packing imparted by the oleic acid moieties. The enhanced fluorescence observed in aqueous media is consistent with aggregation-induced emission (AIE) phenomena reported for certain naphthalimide derivatives, likely arising from restriction of intramolecular motion upon probe aggregation. Confocal fluorescence microscopy of DONDI probes in phosphate-buffered saline (PBS) revealed abundant micron-scale, vesicle-like fluorescent structures, with predominantly spherical particles approximately 1 μm in diameter (data not shown). The solutions visually resembled oil-in-water emulsions, consistent with the presence of oleic acid ester motifs in an aqueous environment. These observations support spontaneous probe self-assembly driven by amphiphilicity and limited aqueous solubility. Importantly, these self-assembled structures are not intended to model physiological lipid assemblies; rather, they illustrate the intrinsic aggregation behavior of the probes in the absence of organized membrane systems.

To distinguish intrinsic probe self-assembly in aqueous buffer from behavior within membrane environments, DONDI analogs were next examined in phase-separated giant unilamellar vesicles (GUVs). When incorporated during electroformation, all probes readily embedded into phospholipid bilayers and preferentially partitioned into the liquid-disordered (Ld) phase of phase-separated GUVs (Figure 2). This membrane-associated behavior contrasts with the higher-order assemblies observed in PBS and indicates that, within a lipid bilayer context, DONDI probes adopt well-defined membrane-partitioning profiles similar to acylglycerols. Preferential localization to the Ld phase is consistent with the presence of unsaturated acyl chains and supports the ability of the DONDI probes to access and report on dynamic, fluid membrane environments. Collectively, these spectroscopic and biophysical data validate the solvatochromic responsiveness and membrane-partitioning properties of the DONDI probes, establishing a foundation for their application in probing lipid organization and dynamics in model membranes and cells.

**Figure 2.**
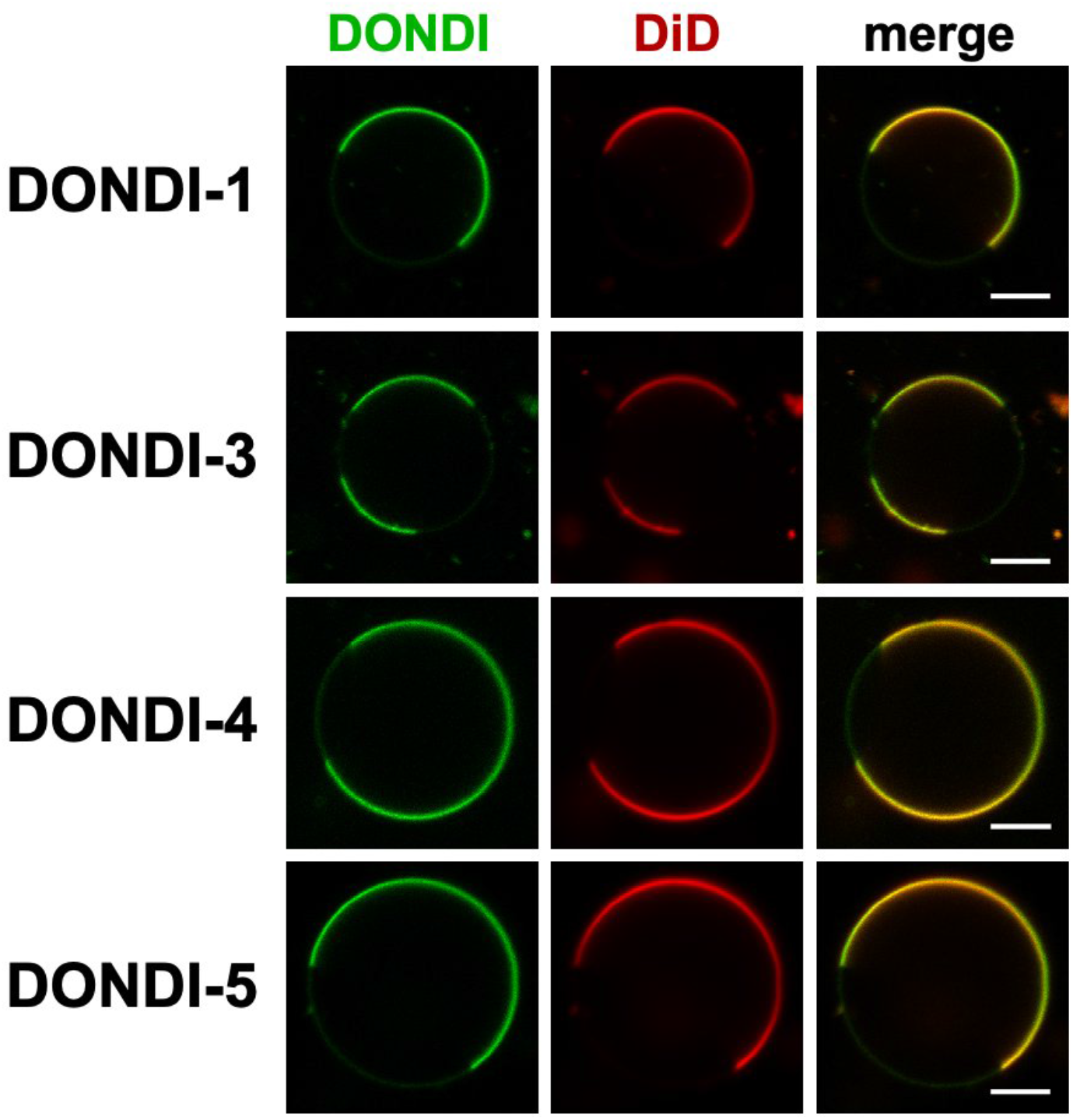
Partitioning of DONDI analogs in phase-separated giant unilamellar vesicles (GUVs). GUVs composed of DOPC/SM/cholesterol (2:2:1 molar ratio) were prepared by electroformation in the presence of DONDI probes (0.1 mol%) and the liquid-disordered (Ld) phase marker DiD. All DONDI analogs preferentially partition into the Ld phase. Scale bar, 10 μm.

### Atomic resolution molecular dynamics simulations

To gain molecular-level insight into the behavior of DONDI probes within lipid bilayers, atomistic molecular dynamics (MD) simulations were performed. Three-dimensional coordinates for all DONDI analogs were constructed using stereochemically defined aminoglycerol scaffolds and parametrized using CHARMM-GUI server.^11–13^ Each simulation system consisted of a POPC bilayer containing a single molecule each of a DONDI probe, diacylglycerol (1,2-dioleoyl-*sn-*glycerol; DAG), triacylglycerol (1,2,3-trioleoyl glycerol; TAG) placed within the same leaflet, while the opposing leaflet contained POPC only. This configuration enabled direct comparison of probe behavior with reference neutral lipids within an identical membrane environment. MD simulations carried out using GROMACS package^14^ revealed both shared and probe-specific features relative to native DAG and TAG. Probe behavior was evaluated by analyzing membrane insertion depth, hydrogen-bonding interactions with lipid headgroups and interfacial water, and the orientation of the naphthalimide fluorophore relative to the membrane plane. Consistent with prior reports for related cholesterol-derived naphthalimide (CND) probes^9^, the chemical identity of the substituent at the C4 position of the naphthalimide core exerted a strong influence on membrane interactions. DONDI-3 and DONDI-4 incorporate polar headgroups (ethanolamine and 4-hydroxypiperidine, respectively) bearing hydroxyl functionality directly linked to the naphthalimide scaffold, whereas DONDI-1 and DONDI-5 contain a more neutral morpholine headgroup. In DONDI-5, fluorophore attachment is shifted from the *sn-*3 to the *sn-*2 position of the glycerol backbone, preserving a free *sn-*3 hydroxyl group and more closely resembling the architecture of native 1,2-DAG. To assess membrane positioning, the distance of the *sn-*3-linked oxygen atom (or the corresponding oxygen atom in DONDI probes) from the membrane center was calculated (Figure 3A). The hydroxyl group distribution of DONDI-3 closely overlapped with that of DAG, indicating a similar average insertion depth. In contrast, DONDI-4 exhibited a broader and slightly shifted distribution, consistent with deeper immersion of the naphthalimide core, in agreement with previous observations for CND probes bearing the same headgroup substitution.^9^ Notably, DONDI-5 displayed a hydroxyl group distribution that closely matched that of 1,2-DAG, consistent with its design to preserve an unmodified *sn-*3 hydroxyl group. Next, we quantified the hydrogen-bonding interactions to assess probe engagement with lipid headgroups and interfacial water. Analysis of the total number of hydrogen bonds (Figure 3B) revealed that DONDI-1 exhibits hydrogen-bonding behavior more closely resembling TAG, whereas DONDI-3, DONDI-4, and DONDI-5 display predominantly DAG-like hydrogen-bonding profiles, albeit with probe-dependent differences. Examination of hydrogen bonds formed specifically with water molecules (Figure 3C) showed that DONDI-3 exhibited the highest degree of water exposure, DONDI-4 the lowest, and DONDI-5 an intermediate profile. These results indicate that while several DONDI analogs retain DAG-like hydrogen-bonding capacity, solvent exposure at the membrane–water interface is strongly modulated by fluorophore placement and hydroxyl accessibility. Because the solvatochromic response of the naphthalimide fluorophore is sensitive not only to membrane insertion depth but also to fluorophore orientation, we next analyzed the tilt angle of the naphthalimide ring relative to the membrane normal (Figure 4). Polar histograms of tilt-angle distributions revealed distinct orientation preferences among the probes. DONDI-3 and DONDI-4 exhibited relatively narrow distributions centered near 90°, indicating preferential alignment of the fluorophore parallel to the membrane plane. In contrast, DONDI-1 showed a broader distribution biased toward smaller tilt angles, consistent with partial alignment along the membrane normal. DONDI-5 showed a pronounced preference for smaller tilt angles relative to the other analogs, indicating a more upright orientation of the naphthalimide ring within the bilayer. Collectively, the MD simulations revealed distinct membrane-interaction profiles among the DONDI probes that directly reflect their molecular design (Figure 5). DONDI-1, DONDI-3, and DONDI-4 share a fully substituted 3-aminoglycerol scaffold bearing two oleic acid chains and exhibit several TAG-like features, particularly in membrane positioning and hydrogen-bonding behavior. In contrast, DONDI-5 contains a single oleic acid chain and a prochiral 2-aminoglycerol scaffold that becomes stereogenic upon asymmetric esterification, preserving a free *sn-*3 hydroxyl group and conferring a more pronounced 1,2-DAG-like character. Together, these features position DONDI-5 as the closest structural and biophysical mimic of native DAG within the DONDI probe series.

**Figure 3.**
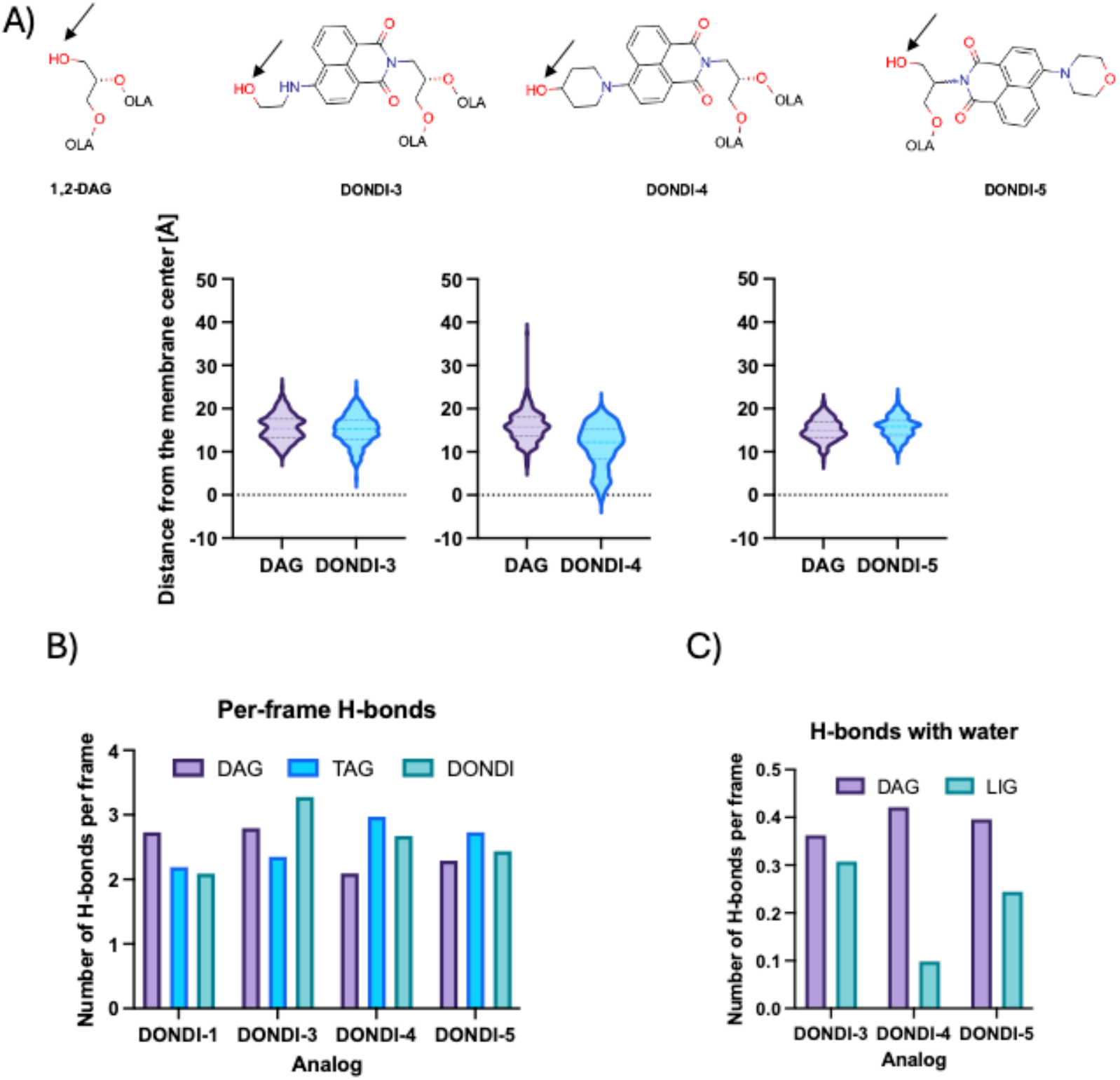
Membrane immersion and hydrogen-bonding analysis of DAG, TAG, and DONDI probes derived from molecular dynamics simulations. **(A)** Position of hydroxyl groups in DONDI analogs relative to the hydroxyl group of 1,2-diacylglycerol (1,2-DAG), quantified as the distance of the hydroxyl oxygen atom (arrow) from the membrane center. **(B)** Total (per-frame) hydrogen-bonding interactions formed by DAG, TAG, and DONDI analogs with surrounding molecules during membrane simulations. **(C)** Hydrogen-bonding interactions of DAG and DONDI analogs specifically with water.

**Figure 4.**
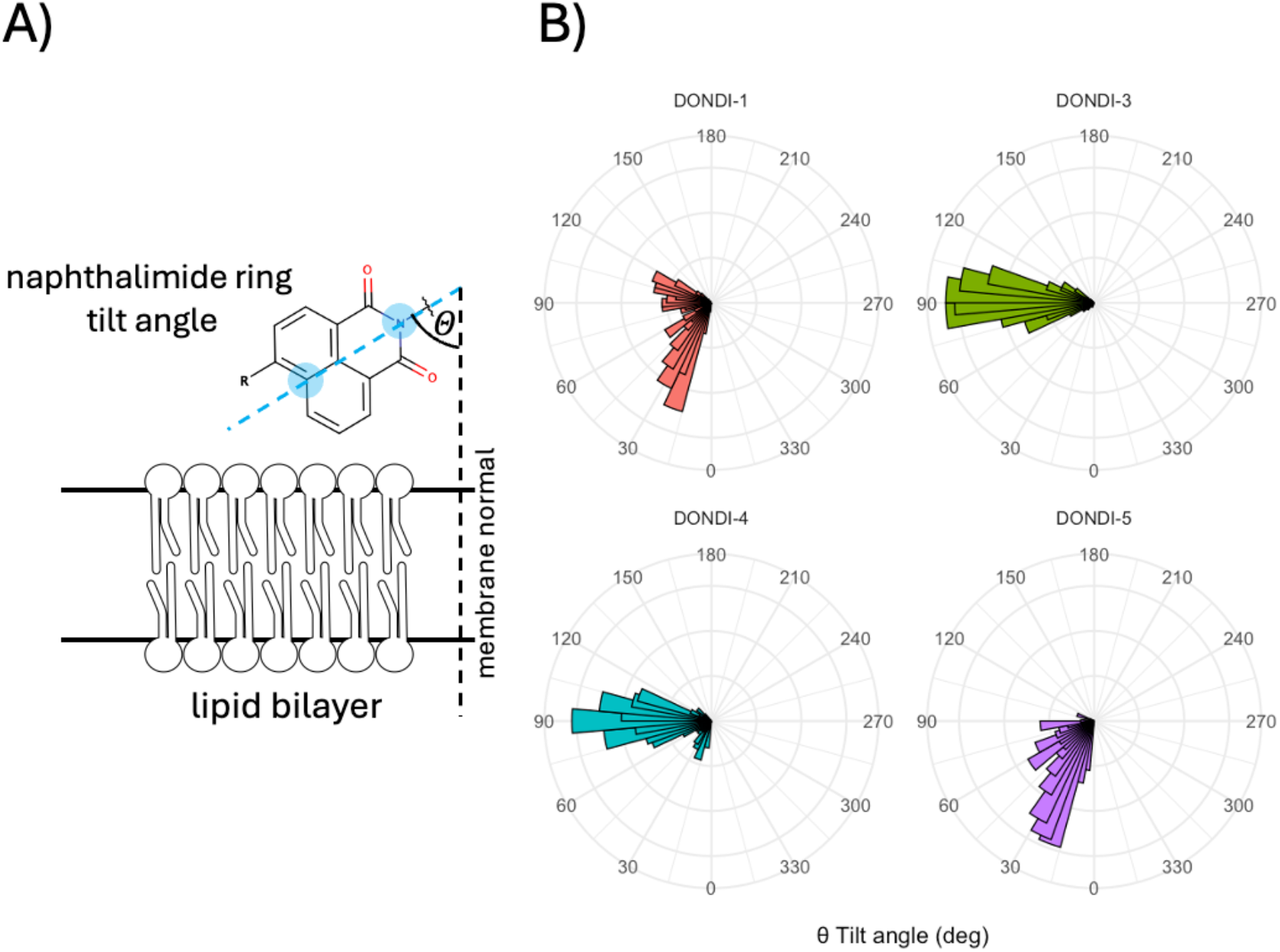
Naphthalimide ring tilt angle analysis in DONDI analogs. **(A)** Atoms of the naphthalimide ring used to define the tilt angle (θ) relative to the membrane normal are highlighted in blue. **(B)** Polar histograms (θ) showing the distribution of tilt angles relative to the membrane normal for the indicated DONDI analogs.

**Figure 5.**
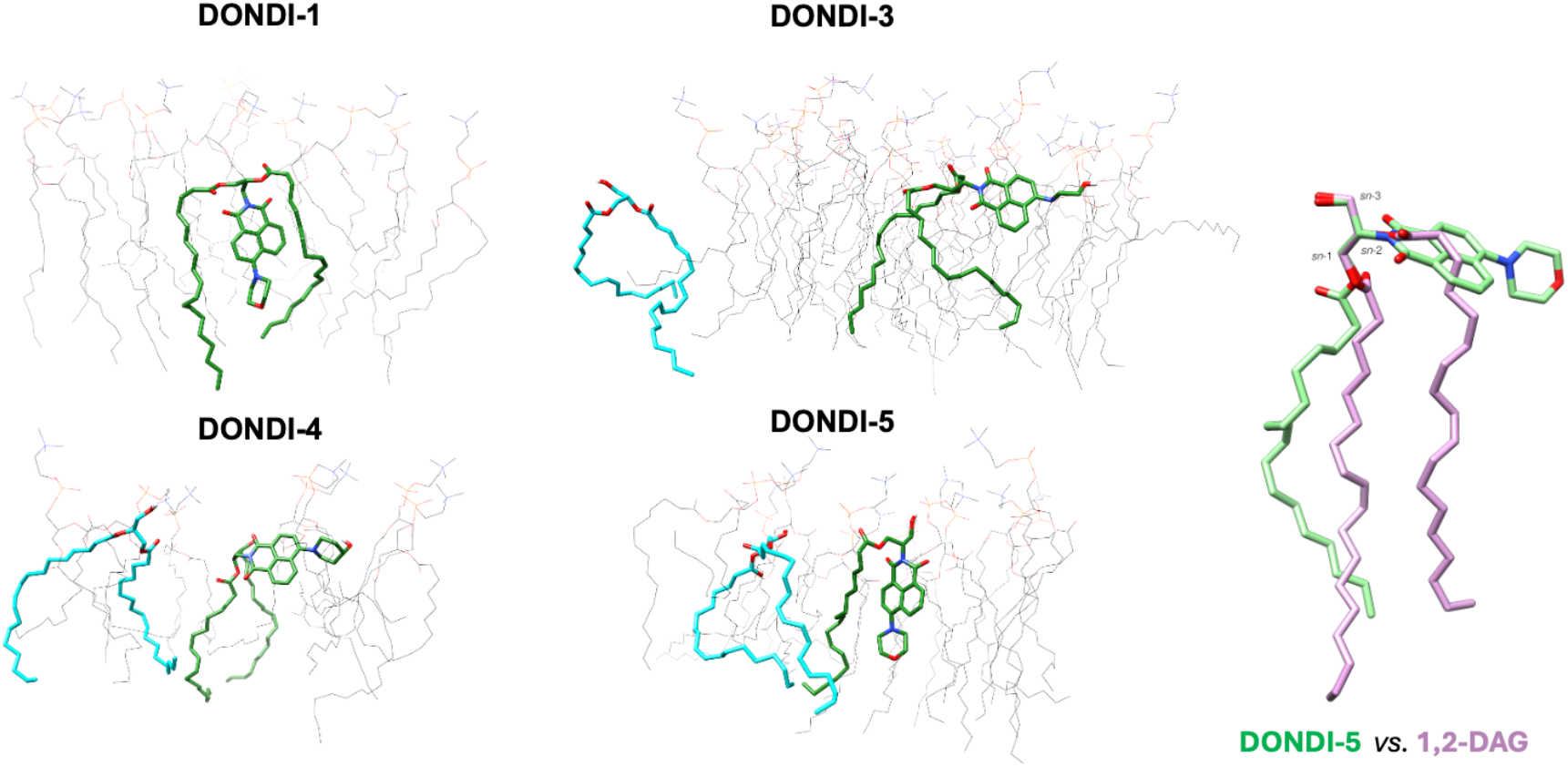
Representative molecular dynamics snapshots illustrating membrane-associated conformations of DONDI probes compared with diacylglycerol (1,2-DAG). DONDI-1, DONDI-3, and DONDI-4 adopt predominantly TAG-like conformations within the bilayer, whereas DONDI-5 exhibits a distinct DAG-like orientation. In DONDI-3 and DONDI-4, a hydroxyl group on the naphthalimide (ND) scaffold promotes a more surface-oriented configuration. In DONDI-5, ND substitution at the *sn*-2 position preserves a free *sn*-3 hydroxyl group, resulting in closer structural alignment with native 1,2-DAG. An overlay of the glycerol scaffolds highlighting the structural similarity between DONDI-5 and 1,2-DAG is shown (right).

### In vitro lipid droplet imaging

Guided by the distinct membrane-interaction profiles revealed by biophysical measurements and molecular dynamics simulations, live-cell imaging of the DONDI analogs was performed using NIH 3T3 fibroblasts. In these experiments, lipid droplet (LD) formation was not stimulated with oleic acid; instead, endogenous LDs were imaged. Probes were introduced into the culture medium from DMSO stock solutions to a final concentration of 1 μM. Cells were incubated with the probes for either 1 h or 24 h prior to imaging. In pulse–chase experiments, cells were incubated with the probes for 1 h, washed with PBS to remove excess probe, and imaged after an additional 24 h. Lipid droplets were visualized using BODIPY 493/503, and LysoView640 was used for lysosomes. Consistent with its predicted DAG-like membrane behavior, the results reveal a clear distinction between DONDI-5 and the other analogs (Figure 6). DONDI-5 rapidly localizes to lipid droplets within 1 h, in contrast to the more TAG-like DONDI analogs, as evidenced by Pearson’s correlation coefficient (PCC) analysis (Figure 6B). The TAG-like analogs exhibit slower uptake kinetics, reaching colocalization levels comparable to DONDI-5 only after 24 h. A similar trend is observed in the pulse–chase experiments, where DONDI-5 stands out with a PCC of approximately 0.7. In addition to strong colocalization with lipid droplets, DONDI5 produces a markedly stronger fluorescence signal than the other analogs, suggesting enhanced accumulation within lipid droplets (Figure 6C). These data demonstrate that DONDI5 exhibits faster and more efficient lipid droplet localization than other TAG-like DONDI analogs. Interestingly, none of these analogs were found to colocalize with lysosomes (PCC< 0.1∼0.2, Figure S2).

**Figure 6.**
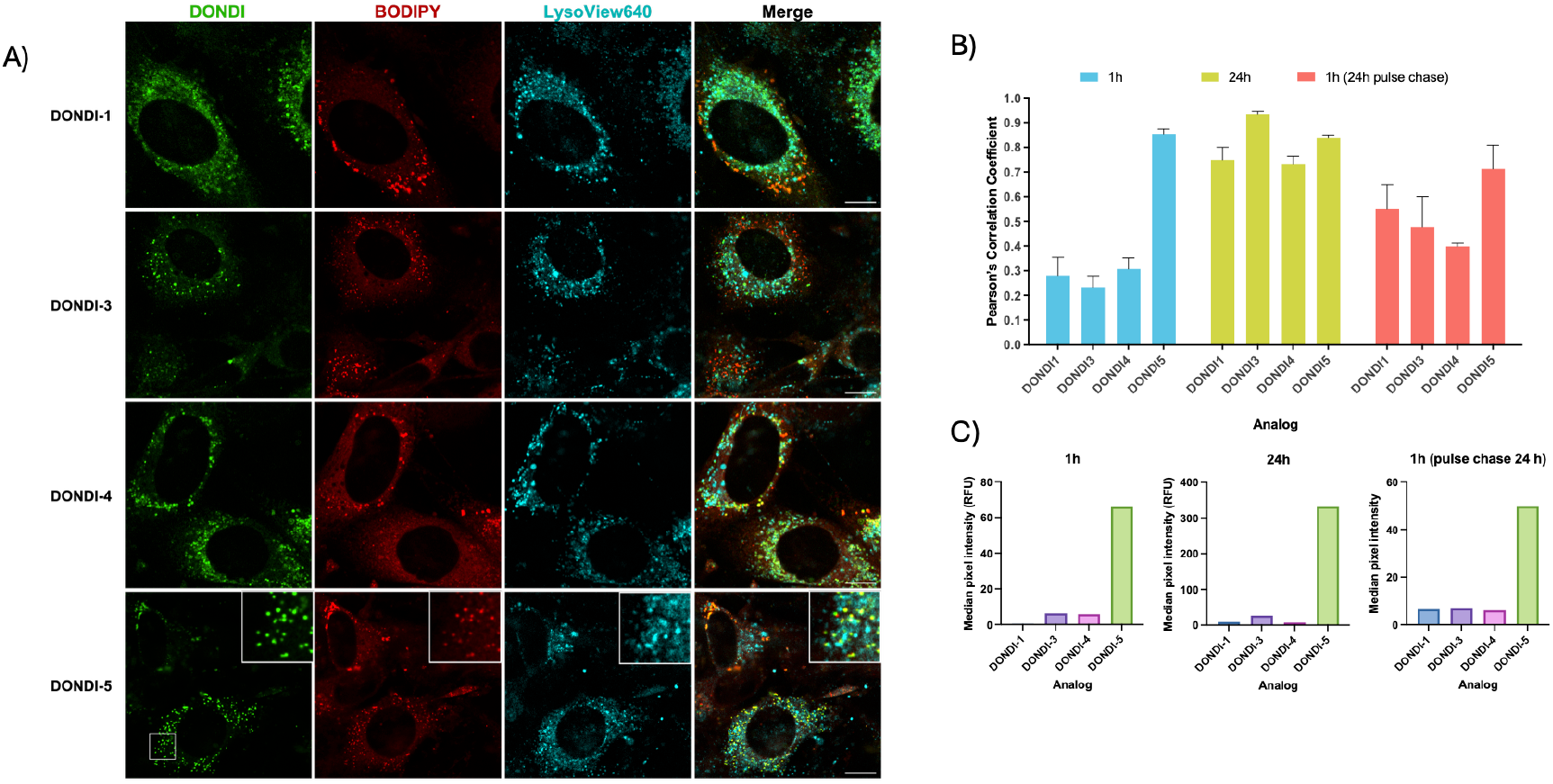
Live-cell imaging and quantitative analysis of DONDI probe localization in NIH 3T3 fibroblasts. **(A)** Confocal fluorescence microscopy images of 3T3 cells loaded with DONDI-1, DONDI-3, DONDI-4, and DONDI-5 (green) after 1h incubation, co-stained with BODIPY 493/503 (red, lipid droplets) and LysoView 640 (cyan, lysosomes). Scale bar, 10 µm. Insets show magnified views of lipid droplets, highlighting probe colocalization. **(B)** Quantification of colocalization between DONDI probes and BODIPY 493/503, expressed as Pearson’s correlation coefficients (PCC), shown as mean ± SD. **(C)** Quantification of DONDI probe fluorescence intensity associated with small intracellular puncta under the same conditions as in panel B. DONDI-5 exhibits rapid lipid droplet enrichment and sustained retention compared to the other analogs.

The pronounced differences in uptake kinetics and fluorescence intensity prompted a closer examination of the structural and metabolic factors that may govern lipid droplet targeting. Lipid droplets (LDs) function as central metabolic hubs, coordinating the trafficking of lipids between organelles to fulfill cellular energy requirements and to provide fatty acid-derived substrates for the biosynthesis of membrane and signaling lipids.^2^ Lipid droplet (LD) biogenesis is a lipid-driven process, with proteins providing supporting and regulatory roles.^15^ With seipin playing a central role in lipid droplet formation, LD growth is initiated by the accumulation of triacylglycerol (TAG) within the endoplasmic reticulum (ER) membrane, leading to the formation of a lipid lens.^16^ TAG biosynthesis begins with glycerol-3-phosphate, which is sequentially converted to diacylglycerol. DAG is then acylated to TAG by diacylglycerol acyltransferase (DGAT) enzymes.^2^ Thus, DAG is an essential substrate for LD formation,^17^ and the development of fluorescent DAG-like probes is of great interest for LD biology. Several fluorescent probes based on DAG and TAG analogs have been reported in the literature: the NBD-aminohexanoic acid conjugate^18,19^, diazo-based photoswitchable fatty acids^20^, BODIPY-tagged DAG^21^, or photocaged DAG analogs.^22^ Small organic fluorescent probes tend to accumulate in lipid droplets and are also used for LDs labeling in cells. Several recent literature reviews summarize their properties and applications.^23–25^ Among small organic fluorescent probes, BODIPY 493/503^26^ has been widely used for live-cell imaging of LDs owing to the well-established photophysical properties of the BODIPY scaffold.^27^ More broadly, small-molecule organic fluorophores offer several advantages, including high cell permeability and readily tunable photophysical characteristics. However, many of these probes display limited specificity for lipid droplets, as they can nonspecifically partition into diverse hydrophobic microenvironments throughout the cell. Even when initial enrichment within lipid droplets is observed, the probes often undergo time-dependent redistribution or diffuse away from the droplets, thereby compromising spatial fidelity. Consequently, their application requires rigorous control over probe incubation and imaging time points to obtain reliable spatial and temporal colocalization data. In contrast, conjugation of small organic fluorophores to native lipid moieties yields probes with greater biological relevance, enabling more accurate monitoring of lipid species localization and intracellular trafficking.^28–30^

Upon addition of DONDI analogs from DMSO solution to the cell culture, they are expected to rapidly interact with the plasma membrane and incorporate into the lipid bilayer. Previous studies on modified diacylglycerol (DAG) analogs added to cell culture have shown rapid trans bilayer movement across the plasma membrane following cellular uptake, suggesting that a similar mechanism may operate for the DONDI probes.^31^ On this basis, we hypothesize that the initial membrane insertion and subsequent intracellular trafficking of our analogs occur on a rapid timescale. Several potential metabolic fates for exogenous DAG-like molecules have been described. Florin-Christensen *et al*. demonstrated that radioactively labeled 1,2-dioctanoylglycerol introduced into multiple cell lines, including NIH 3T3 fibroblasts, underwent direct conversion to dioctanoylphosphatidylcholine and dioctanoyltriacylglycerol, as well as lipolytic degradation. In contrast, 1-oleoyl-2-acetyl-*sn*-glycerol was metabolized exclusively via lipolysis, highlighting how subtle structural differences can dramatically alter intracellular processing.^32^ This study also proposed the existence of subcellular mechanisms for DAG transport. Consistent with these findings, subsequent work confirmed the hydrolysis of exogenously supplied DAG species, such as 1-palmitoyl-2-oleoyl-*sn*-glycerol and 1-stearoyl-2-arachidonoyl-*sn*-glycerol, in smooth muscle cells *via* DAG lipase pathways, yielding primarily monoacylglycerols and free fatty acids, with minor formation of triacylglycerols and phosphatidylcholines.^33^ Together, these studies underscore the complexity of intracellular lipid handling pathways and emphasize that the metabolic fate of exogenous DAGs is highly structure-dependent. Ho and Storch demonstrated that Caco-2 cells exhibit saturable uptake of *sn*-2-monoolein at both the apical and basolateral membranes, consistent with protein-mediated transport involving plasma membrane fatty acid-binding protein (FABP_pm_) and fatty acid transport proteins rather than passive diffusion.^34^ Competition between monoacylglycerols and long-chain fatty acids suggests shared carrier pathways.^35^ Further evidence shows that long-chain fatty acids and monoacylglycerols are transported into adipocytes (3T3-L1) by fatty acid transport proteins 1 and 4.^36^ Additional studies further support a direct connection between DAG metabolism and LD growth. Incubation of hepatoma cells with monoacylglycerols was shown to trigger DGAT2 translocation to LDs, promoting LD expansion through coordinated activity with monoacylglycerol acyltransferase (MGAT2).^37^ Similarly, treatment with 2-acylglycerols induced LD formation comparable to that induced by oleic acid supplementation, reinforcing the role of DAG-derived intermediates in neutral lipid storage.

The LD themselves have been shown to contain significant pools of DAGs. Thiele et al. reported that DAGs present in lipid droplets of cultured adipocytes and fibroblasts are dynamically esterified to triacylglycerols via LD-associated DGAT2.^38^ Notably, when triglyceride synthesis was inhibited, fluorescent DAG analogs accumulated in the endoplasmic reticulum, yet substantial amounts still entered lipid droplets *via* an as-yet-unknown mechanism. These observations point to previously unrecognized pathways that promote DAG enrichment in LDs without prior conversion to triacylglycerols.

Motivated by fluorescence imaging of DONDI-5, we sought to determine the chemical identity of the fluorescent puncta observed within lipid droplets. To address this question, cells were incubated with DONDI-5 (10 µM, to improve subsequent analysis) for 24 h, after which culture media and cell pellets were collected separately. Lipids were extracted from both fractions and analyzed by reverse-phase HPLC equipped with a fluorescence detector to identify DONDI-5 and its metabolites. As shown in Figure 7A, reverse-phase HPLC analysis revealed a dominant peak at 3.80 min corresponding to intact DONDI-5, which was detected at much higher levels in the cell pellet. Quantitative analysis indicated that ∼ 70% of intracellular DONDI-5 remained intact after 24 h of incubation (Figure 7B). A minor peak at 1.14 min was assigned to 2-(naphthalimido)glycerol, consistent with a lipolytic hydrolysis product. In contrast, analysis of the culture medium revealed only 4.5% of intact DONDI-5, with 95.5% of the signal corresponding to aminoglycerol-naphthalimide, consistent with extracellular hydrolysis and/or release of the hydrolyzed product.

**Figure 7.**
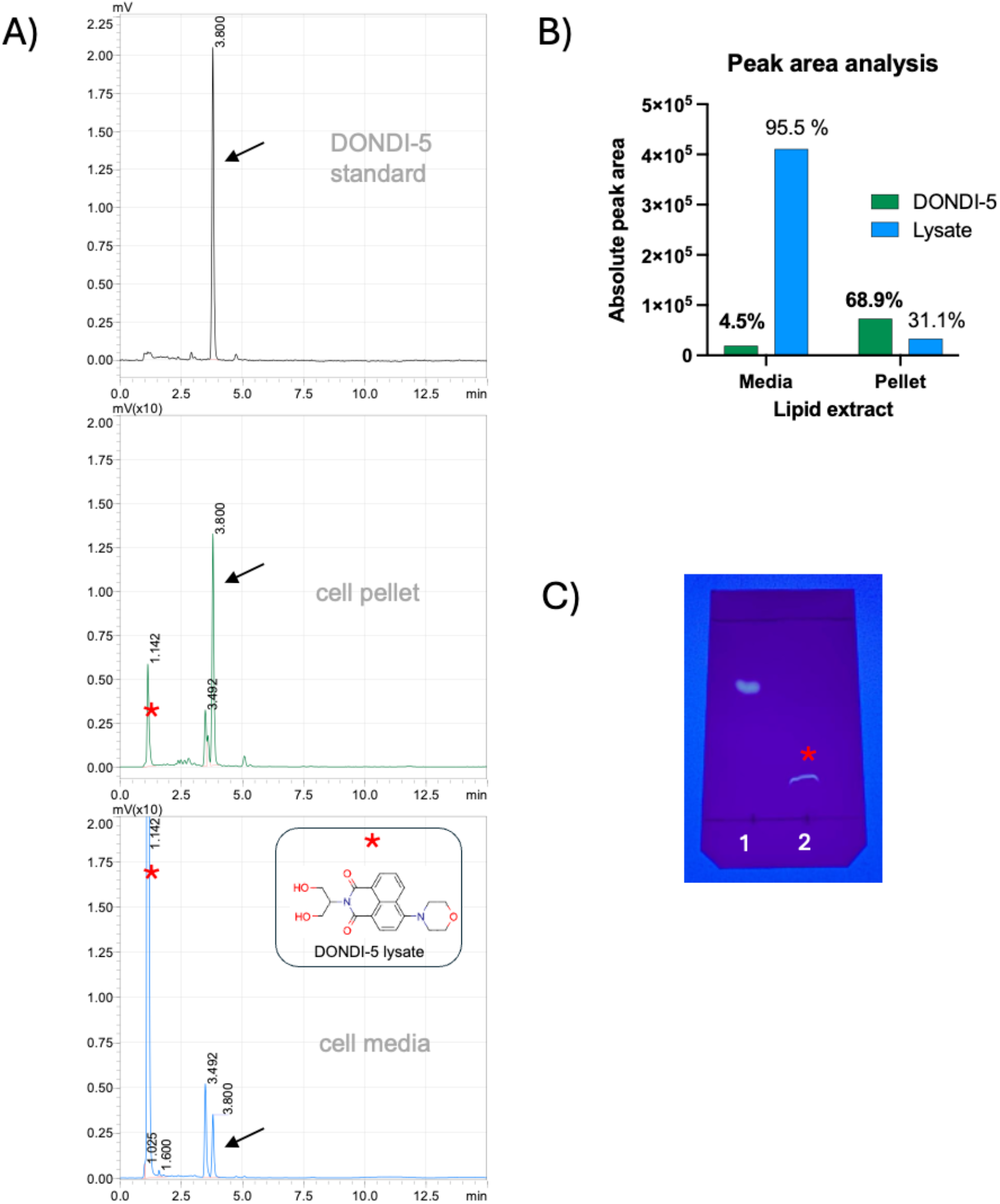
Reverse-phase HPLC and TLC analysis of DONDI-5 and its metabolites following cellular uptake. **(A)** Representative fluorescence-detected RP-HPLC chromatograms of lipid extracts from cell pellets and culture medium, compared with a DONDI-5 standard. The peak at 3.80 min (arrow) corresponds to intact DONDI-5, whereas the peak at 1.14 min (red asterisk) corresponds to the hydrolysis product, 2-(naphthalimido)glycerol. **(B)** Quantitative peak area analysis of intact and hydrolyzed DONDI-5 in lipid extracts from culture medium and cell pellets, expressed as percentages of total DONDI-5-derived signal. **(C)** Thin-layer chromatography (TLC) analysis of DONDI-5 incubated with porcine cholesteryl esterase and visualized under UV illumination (395 nm), demonstrating enzymatic hydrolysis. Lane 1, DONDI-5 standard; lane 2, DONDI-5 after esterase treatment.

To further confirm the identity of the DONDI-5 hydrolysis product, DONDI-5 was incubated with a readily accessible model lipase, porcine cholesteryl esterase (CEase), followed by TLC and HPLC analysis (Figure 7). Cholesteryl esterase (EC 3.1.1.13), also known as bile salt– activated lipase or carboxyl ester lipase, is a broadly acting neutral lipid hydrolase capable of cleaving cholesteryl esters, triacylglycerols, phospholipids, and other hydrophobic ester substrates.^39,40^ Owing to its broad substrate specificity, CEase was employed here as a generic lipase to assess ester bond accessibility within the DONDI scaffold. This treatment resulted in the quantitative formation of 2-(naphthalimido)glycerol, which had the identical retention time (1.14 min) as the hydrolysis product detected in cellular extracts. Consistent with this rationale, all DONDI analogs were found to be efficient substrates for CEase-mediated hydrolysis. Taken together, the HPLC data indicate that DONDI-5, localized to LDs, remains largely intact over prolonged incubation, whereas degradation products are released from cells.

## CONCLUSIONS

Collectively, our studies establish DONDI probes as environment-sensitive, membrane-partitioning fluorescent reporters whose behavior is governed by both fluorophore solvatochromism and lipid-mimetic molecular design. Spectroscopic analyses confirmed pronounced solvatochromic responses and large Stokes shifts characteristic of naphthalimide fluorophores, while biophysical experiments revealed efficient incorporation into model membranes and liquid-disordered lipid phases. Atomic-resolution molecular dynamics simulations further demonstrated that subtle differences in glycerol substitution and head-group chemistry strongly influence membrane insertion depth, hydrogen-bonding patterns, and fluorophore orientation, with DONDI-5 most closely recapitulating key structural and interfacial features of 1,2-diacylglycerol. Consistent with these properties, live-cell imaging revealed rapid and sustained localization of DONDI-5 to LDs, distinguishing it from more TAG-like analogs, and biochemical analysis confirmed that the probe remains largely intact following intracellular accumulation. Taken together, DONDI-5 represents a chemically stable, diacylglycerol-mimetic fluorescent probe that enables visualization of DAG-associated LD dynamics and acylglycerol partitioning in living cells, providing a broadly applicable chemical tool for probing lipid metabolic pathways and ER-LD axis with spatial and temporal resolution. The internalization of DONDI-5 and its accumulation in lipid droplets support the possibility of lipid trafficking pathways that enable delivery to lipid droplets independently of canonical DAG-to-TAG conversion. Ongoing studies aim to directly test this model and to define the molecular determinants underlying this alternative trafficking route.

## ASSOCIATED CONTENT

### Supporting Information

The Supporting Information is available free of charge at https://pubs.acs.org/doi/

Additional experimental details, including synthetic procedures, compound characterization data, molecular dynamics simulation protocols, and other experimental methods.

## AUTHOR INFORMATION

### Notes

The authors declare no competing financial interest.

## ACKNOWLEDGMENTS

This work was supported by the Florida Atlantic University Stiles–Nicholson Brain Institute seed grant and, in part, by National Institutes of Health grant 1R15GM147912-01, which supported HPLC instrumentation and, in part, by National Institutes of Health grant 1R15AG085620-01, which supported cell culture. We gratefully acknowledge the Florida Atlantic University Office of Undergraduate Research and Inquiry for continued support of undergraduate research.

