## Supplementary_information for "Chemical Probing of Diacylglycerol Dynamics at Lipid Droplets"

### Table of Contents

| | | PAGE |
|---|---|---|
| | **Supplementary figures and tables referenced in the main text** | S3-S4 |
| | **Materials and Methods** | |
| | Synthetic methods | S5-S8 |
| | Photophysical characterization of compounds | S9 |
| | Molecular Dynamics Simulations and Analysis | S9-S10 |
| | Cell Culture and Fluorescence Confocal Imaging | S10-S11 |
| | HPLC and TLC Analysis | S11-S12 |
| | NMR and MS spectra of all intermediates and final compounds | S13-21 |
| | **SI References** | S22 |

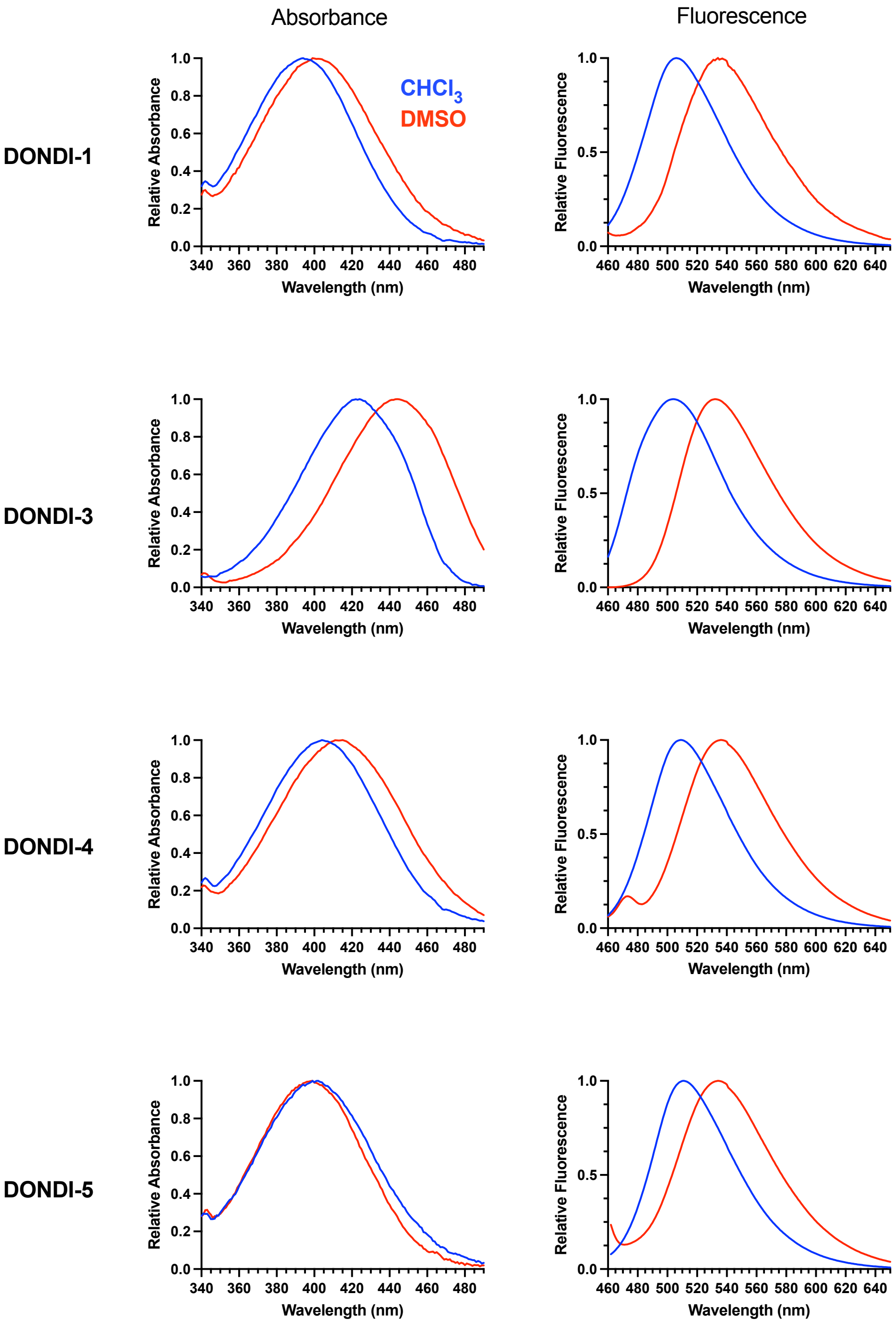

**Figure S1:** Absorbance and fluorescence spectra profiles of DONDI analogs in chloroform and DMSO.

**Table S1.** Excitation and emission data summary from Figure S1:

| | | DONDI-1 | DONDI-3 | DONDI-4 | DONDI-5 |
|---|---|---|---|---|---|
| **Chloroform** | Absorbance ($\lambda_{max}$) | 392 | 424 | 405 | 399 |
| | Emission (at $\lambda_{max}$) | 506 | 503 | 509 | 511 |
| **DMSO** | Absorbance ($\lambda_{max}$) | 399 | 444 | 415 | 402 |
| | Emission (at $\lambda_{max}$) | 540 | 532 | 540 | 540 |

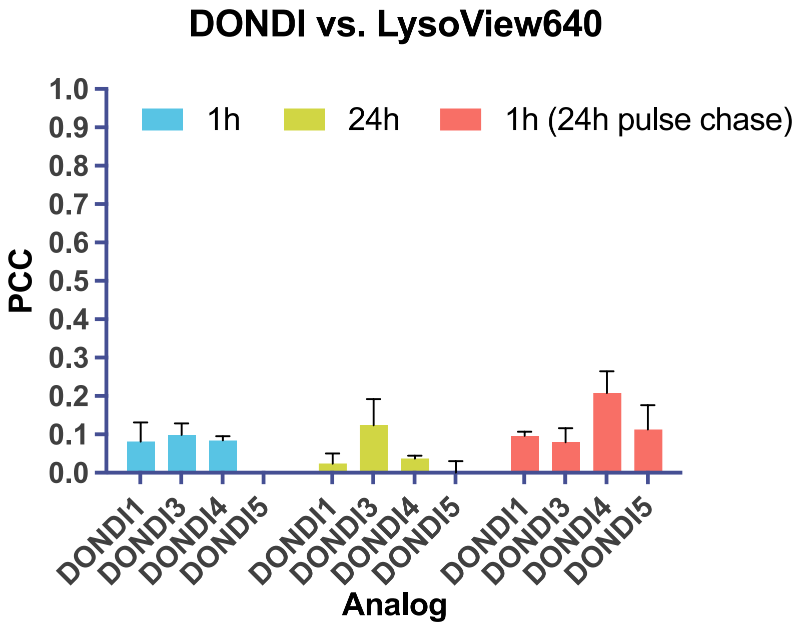

**Figure S2. Pearson's correlation coefficient (PCC) analysis of DONDI probe colocalization with lysosomes over time.** PCC values (mean ± SD) quantify the extent of colocalization between DONDI analogs and the lysosomal marker LysoView 640 in NIH 3T3 fibroblasts at the indicated time points. Across all conditions examined, PCC values remain low, indicating minimal lysosomal localization of all DONDI probes.

### Materials and methods

**Reagents, solvents, and glassware.** All reagents and solvents, including anhydrous solvents, were obtained from commercial suppliers (Fisher Scientific) and used without further purification unless otherwise stated. LysoView™ 640 was purchased from Biotium, and BODIPY 493/503 was obtained from Fisher Scientific.
**Thin-layer chromatography (TLC).** Thin-layer chromatography (TLC) was performed on aluminum-backed silica gel plates (60 Å pore diameter, 200 µm layer thickness; SiliCycle or EMD Millipore). Plates containing a fluorescent indicator were visualized under UV illumination ($\lambda_{ex}$ = 254 or 366 nm) or by staining with 10% $H_2SO_4$ in ethanol, followed by brief heating with a heat gun.
**Column chromatography.** Column chromatography was performed using silica gel (60 Å pore diameter, 40–60 µm particle size; SiliCycle).
**NMR and Mass Spectrometry.** NMR spectra were recorded on a 400 MHz spectrometer at 22 °C in $CDCl_3$ or $CD_3OD$. Proton ($^1H$) and carbon ($^{13}C$) chemical shifts were referenced to tetramethylsilane (TMS). Signal multiplicities are denoted as singlet (s), doublet (d), triplet (t), quartet (q), pentet (p), multiplet (m), broad (br), or combinations thereof. Mass spectra were acquired on a Bruker Microflex LRF MALDI-TOF mass spectrometer using α-cyano-4-hydroxycinnamic acid (α-CHCA) as the matrix and recorded in positive ion mode.

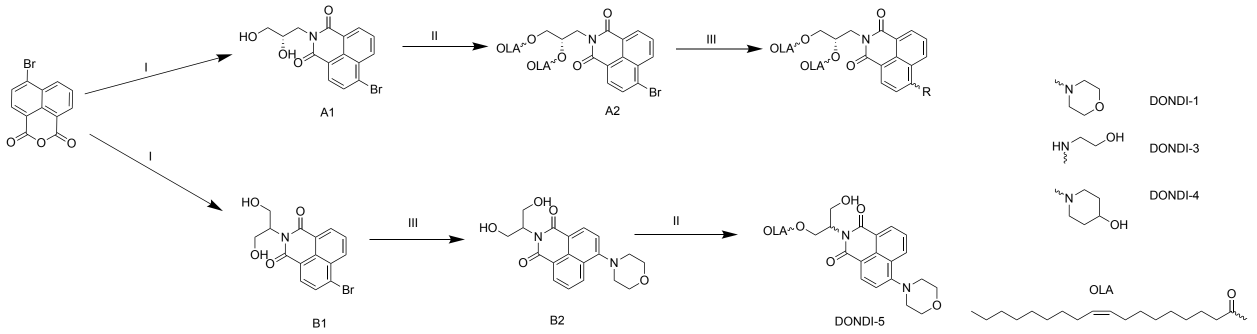

**Scheme1:** Synthesis of DONDI analogs. Reagents and conditions: (I) (S)-3-amino-1,2-propanediol or 2-amino-1,2-propanediol, 4-bromo-1,8-naphthalic anhydride, diisopropylethylamine (DIPEA), ethanol, reflux. (II) Oleic acid, N,N′-dicyclohexylcarbodiimide (DCC), 4-dimethylaminopyridine (DMAP), chloroform. (III) A2 or B1, methoxyethanol, corresponding amine (morpholine, ethanolamine, or 4-hydroxypiperidine), reflux.

#### Synthesis of A1

(S)-3-Amino-1,2-propanediol (4 mmol), 4-bromo-1,8-naphthalic anhydride (4 mmol), and N,N-diisopropylethylamine (DIPEA, 4 mmol) were dissolved in ethanol (25 mL) in a round-bottom flask. The reaction mixture was heated at reflux for 1 h. After completion, the solvent was removed under reduced pressure. The resulting residue was dissolved in ethyl acetate and transferred to a separatory funnel, where it was washed with 10% aqueous HCl. The organic layer was separated,

dried over anhydrous $Na_2SO_4$, filtered, and concentrated under reduced pressure to afford the desired product as a brown solid (71% yield).
$^1$H-NMR (400 MHz, DMSO-$d_6$) δ 8.49 (ddd, J = 16.9, 7.9, 1.1 Hz, 2H), 8.27 (d, J = 7.9 Hz, 1H), 8.16 (d, J = 7.9 Hz, 1H), 7.95 (dd, J = 8.5, 7.3 Hz, 1H), 4.80 (d, J = 5.2 Hz, 1H), 4.62 (t, J = 5.7 Hz, 1H), 4.20 (dd, J = 12.8, 8.2 Hz, 1H).

**Synthesis of A2**

A1 (7.11 mmol) and 4-dimethylaminopyridine (DMAP, 2.2 equiv) were dissolved in anhydrous chloroform (40 mL) and tetrahydrofuran (3 mL) in a round-bottom flask under anhydrous conditions. Oleic acid (2.2 equiv) was added, and the reaction mixture was stirred at room temperature for 15 min. A separate solution of N,N′-dicyclohexylcarbodiimide (DCC, 2.2 equiv) in anhydrous chloroform (5 mL) was then added dropwise. The reaction mixture was cooled in an ice bath for 60 min and subsequently allowed to warm to room temperature and stir overnight. The solvent was removed under reduced pressure, and the residue was dissolved in ethyl acetate and filtered to remove the dicyclohexylurea byproduct. The filtrate was washed with water and brine, dried over anhydrous $Na_2SO_4$, filtered, and concentrated under reduced pressure. The crude product was purified by flash column chromatography on silica gel (5:1 hexanes/ethyl acetate) to afford the desired product as a yellow oil (66% yield).
$^1$H-NMR (400 MHz, $CDCl_3$) δ 8.67 (dd, J = 7.3, 1.2 Hz, 1H), 8.61 (dd, J = 8.5, 1.1 Hz, 1H), 8.43 (d, J = 7.9 Hz, 1H), 8.07 (d, J = 7.9 Hz, 1H), 7.87 (dd, J = 8.5, 7.3 Hz, 1H), 5.55 (ddt, J = 7.7, 5.8, 3.8 Hz, 1H), 5.42 – 5.27 (m, 4H), 4.64 (dd, J = 13.7, 7.8 Hz, 1H), 4.44 – 4.23 (m, 3H), 2.35 (t, J = 7.6 Hz, 2H), 2.23 (t, J = 7.5 Hz, 2H), 2.07 – 1.97 (m, 7H), 2.00 – 1.93 (m, 1H), 1.68 – 1.58 (m, 2H), 1.45 (td, J = 12.5, 5.5 Hz, 2H), 1.38 – 1.22 (m, 37H), 1.14 (q, J = 4.0 Hz, 6H), 0.96 – 0.85 (m, 5H). MALDI-TOF: $[M+Na]^+$ calcd. m/z = 902.4736, obs. m/z = 902.2990

**Synthesis of DONDI-1**

A2 (1.323 mmol) was dissolved in methoxyethanol (7 mL) in a round-bottom flask and stirred. Morpholine (5 equiv) was added, and the reaction mixture was heated at reflux overnight. After cooling to room temperature, the reaction mixture was diluted with water and brine and acidified with 10% (v/v) aqueous HCl. The aqueous phase was extracted with dichloromethane (3 × 25 mL). The combined organic extracts were dried over anhydrous $Na_2SO_4$, filtered, and concentrated under reduced pressure. The crude product was purified by flash chromatography on a Büchi Pure C-810 system using a 12 g EcoFlex silica gel column (5% ethyl acetate in dichloromethane, 10 min isocratic) to afford the desired product as a yellow solid (42% yield).
$^1$H-NMR (400 MHz, $CDCl_3$) δ 8.58 (dd, J = 7.3, 1.2 Hz, 1H), 8.53 (d, J = 8.1 Hz, 1H), 8.43 (dd, J = 8.5, 1.2 Hz, 1H), 7.70 (dd, J = 8.5, 7.3 Hz, 1H), 7.23 (d, J = 8.1 Hz, 1H), 5.54 (ddt, J = 7.7, 5.9, 3.9 Hz, 1H), 5.40 – 5.32 (m, 1H), 5.36 – 5.25 (m, 3H), 4.61 (dd, J = 13.6, 7.6 Hz, 1H), 4.42 – 4.21 (m, 3H), 4.02 (dd, J = 5.7, 3.4 Hz, 4H), 3.30 – 3.23 (m, 4H), 2.33 (t, J = 7.6 Hz, 2H), 2.22 (t, J = 7.5 Hz, 2H), 2.07 – 1.92 (m, 8H), 1.62 (s, 1H), 1.61 (d, J = 13.7 Hz, 2H), 1.45 (ddd, J = 13.7, 9.5, 6.3 Hz, 2H), 1.33 (d, J = 7.4 Hz, 2H), 1.31 – 1.21 (m, 34H), 1.15 (q, J = 3.9 Hz, 6H), 0.91 – 0.84 (m, 5H). MALDI-TOF: $[M+Na]^+$ calcd. m/z = 907.6171, obs. m/z = 907.6956

**Synthesis of DONDI-3**

A2 (0.4 mmol) was dissolved in methoxyethanol (7 mL) in a round-bottom flask and stirred. Ethanolamine (5 equiv) was added, and the reaction mixture was heated at reflux overnight. After cooling to room temperature, the mixture was diluted with water and brine and acidified with 10% (v/v) aqueous HCl. The aqueous phase was extracted with dichloromethane (3 × 25 mL). The combined organic extracts were dried over anhydrous $Na_2SO_4$, filtered, and concentrated under reduced pressure. The crude product was purified by flash column chromatography on silica gel using a gradient of hexanes/ethyl acetate (1:1 followed by 2:1) to afford the desired product as a yellow solid (84% yield).
$^1$H-NMR (400 MHz, $CDCl_3$) δ 8.44 – 8.33 (m, 2H), 8.03 (dd, J = 8.6, 1.1 Hz, 1H), 7.42 (dd, J = 8.4, 7.3 Hz, 1H), 6.64 (d, J = 8.5 Hz, 1H), 5.89 (t, J = 5.2 Hz, 1H), 5.57 (tt, J = 6.7, 4.0 Hz, 1H), 5.33 (td, J = 6.1, 3.6 Hz, 4H), 4.54 (dd, J = 13.6, 7.2 Hz, 1H), 4.43 (dd, J = 12.0, 3.6 Hz, 1H), 4.28 (ddd, J = 15.7, 12.8, 5.4 Hz, 2H), 4.04 (q, J = 5.2 Hz, 2H), 3.54 (q, J = 5.1 Hz, 2H), 2.44 (t, J = 5.5 Hz, 1H), 2.30 (dt, J = 22.6, 7.5 Hz, 4H), 2.04 – 1.92 (m, 8H), 1.66 – 1.55 (m, 3H), 1.50 (tt, J = 9.8, 5.1 Hz, 2H), 1.37 – 1.24 (m, 35H), 1.21 – 1.14 (m, 6H), 0.91 – 0.83 (m, 5H). MALDI-TOF: $[M+Na]^+$ calcd. m/z = 881.6014, obs. m/z = 881.6268

**Synthesis of DONDI-4**

A2 (0.3 mmol) was dissolved in methoxyethanol (7 mL) in a round-bottom flask and stirred. 4-hydroxypiperidine (5 equiv) was added, and the reaction mixture was heated at reflux overnight. After cooling to room temperature, the reaction mixture was diluted with water and brine and acidified with 10% (v/v) aqueous HCl. The aqueous phase was extracted with dichloromethane (3 × 25 mL). The combined organic extracts were dried over anhydrous $Na_2SO_4$, filtered, and concentrated under reduced pressure. The crude product was purified by flash column chromatography using silica gel using dichloromethane/ethyl acetate (10:1) to afford the desired product as a yellow solid (44% yield).
$^1$H-NMR (400 MHz, $CDCl_3$) δ 8.57 (dd, J = 7.3, 1.2 Hz, 1H), 8.49 (d, J = 8.1 Hz, 1H), 8.38 (dd, J = 8.5, 1.3 Hz, 1H), 7.69 (dd, J = 8.4, 7.3 Hz, 1H), 7.20 (d, J = 8.1 Hz, 1H), 5.54 (ddt, J = 7.7, 6.0, 3.9 Hz, 1H), 5.40 – 5.25 (m, 4H), 4.60 (dd, J = 13.7, 7.6 Hz, 1H), 4.42 – 4.21 (m, 3H), 4.02 (dq, J = 8.6, 4.3 Hz, 1H), 3.53 (dt, J = 11.4, 4.8 Hz, 2H), 3.08 (ddd, J = 12.3, 9.2, 2.9 Hz, 2H), 2.32 (t, J = 7.6 Hz, 2H), 2.20 (dt, J = 17.4, 8.8 Hz, 4H), 1.96 (ttd, J = 17.8, 7.5, 3.9 Hz, 10H), 1.70 – 1.55 (m, 3H), 1.48 (s, 2H), 1.46 (dt, J = 7.0, 3.4 Hz, 1H), 1.36 – 1.19 (m, 36H), 1.19 – 1.11 (m, 2H), 1.15 (s, 4H), 0.91 – 0.83 (m, 6H). MALDI-TOF: $[M+H]^+$ calcd. m/z = 899.6508, obs. m/z = 899.6762

**Synthesis of B1**

(S)-2-Amino-1,2-propanediol (4 mmol, 2 equiv) and 4-bromo-1,8-naphthalic anhydride (2 mmol, 1 equiv) were dissolved in ethanol (10 mL) in a round-bottom flask. The reaction mixture was heated at reflux for 4 h. After completion, the solvent was removed under reduced pressure, and the resulting precipitate was suspended in ethyl acetate and collected by filtration. The solid was dried in an oven at 110 °C to afford the desired product as a brown solid (71% yield).
$^1$H-NMR (400 MHz, DMSO-$d_6$) δ 8.47 (d, J = 7.2 Hz, 1H), 8.40 (dd, J = 8.5, 1.1 Hz, 1H), 8.23 (d, J = 7.9 Hz, 1H), 8.11 (d, J = 7.9 Hz, 1H), 7.90 (dd, J = 8.5, 7.3 Hz, 1H), 5.15 (tt, J = 8.0, 5.8 Hz, 1H), 4.75 (s, 2H), 3.90 (t, J = 9.7 Hz, 2H), 3.75 (dt, J = 10.6, 4.6 Hz, 2H).

**Synthesis of B2**

B1 was dissolved in DMSO (5 mL) in a round-bottom flask and stirred. Morpholine (5 equiv) was added, and the reaction mixture was heated at reflux overnight. After cooling to room temperature, the mixture was diluted with water and brine and acidified to pH 6–7 with 1% aqueous HCl. The aqueous phase was extracted with chloroform (3 × 10 mL). The combined organic layers were dried over anhydrous $Na_2SO_4$, filtered, and concentrated under reduced pressure to afford a yellow solid. The crude material was dry-loaded onto silica gel and purified by flash column chromatography (chloroform/methanol, 10:1) to yield the desired product as a light yellow solid (55% yield).

**Synthesis of DONDI-5**

B2 (1 mmol) and 4-dimethylaminopyridine (DMAP, 0.1 equiv) were dissolved in anhydrous chloroform (40 mL) and tetrahydrofuran (3 mL) under anhydrous conditions in a round-bottom flask. Oleic acid (1.1 equiv) was added, and the reaction mixture was stirred at room temperature for 15 min. A solution of N,N′-dicyclohexylcarbodiimide (DCC, 1.1 equiv) in anhydrous chloroform (5 mL) was added dropwise, and the reaction mixture was cooled in an ice bath for 60 min before being allowed to warm to room temperature and stirred overnight. The solvent was removed under reduced pressure, and the residue was dissolved in ethyl acetate and filtered to remove the dicyclohexylurea byproduct. The filtrate was washed sequentially with water, brine, and aqueous HCl, and the aqueous phase was extracted with chloroform (3 × 25 mL). The combined organic extracts were dried over anhydrous $Na_2SO_4$, filtered, and concentrated under reduced pressure to afford a yellow oil. Purification by flash column chromatography on silica gel (chloroform/ethyl acetate, 1:1) afforded the desired product as a yellow solid (10% yield).
$^1$H-NMR (400 MHz, $CDCl_3$) δ 8.60 (d, J = 7.3 Hz, 1H), 8.54 (d, J = 8.0 Hz, 1H), 8.45 (dd, J = 8.4, 1.2 Hz, 1H), 7.73 (dd, J = 8.5, 7.3 Hz, 1H), 7.26 (d, J = 8.1 Hz, 1H), 5.66 (tt, J = 7.4, 5.9 Hz, 1H), 5.34 (q, J = 5.9 Hz, 2H), 4.77 – 4.61 (m, 2H), 4.04 (dd, J = 5.8, 3.3 Hz, 2H), 3.32 – 3.26 (m, 2H), 2.25 (t, J = 7.5 Hz, 2H), 2.00 (dq, J = 13.3, 6.7 Hz, 5H), 1.53 (p, J = 7.4 Hz, 3H), 1.38 – 1.22 (m, 19H), 1.19 (t, J = 5.2 Hz, 7H), 0.95 – 0.85 (m, 4H). MALDI-TOF: $[M+Na]^+$ calcd. m/z = 643.3718, obs. m/z = 642.4801

**Determination of Absorbance and Fluorescence Spectra in Organic Solvents**

Stock solutions of DONDI probes were diluted to 10 μM in chloroform and evaporated to dryness. The resulting residues were re-dissolved in chloroform or dimethyl sulfoxide (DMSO) to afford 10 μM solutions. Absorbance spectra were recorded from 340 to 490 nm using a Jasco FP-8550 spectrofluorometer operated in absorbance mode. Fluorescence emission spectra were acquired in the corresponding solvents using fluorescence mode, with excitation at the absorbance maximum ($\lambda_{ex} = \lambda_{max}$), and recorded over the range of 460–650 nm.

**Visualization of Lipid Phase Partitioning in Giant Unilamellar Vesicles (GUVs)**

Lipid stock solutions of DOPC (10 mM), brain sphingomyelin (10 mM), cholesterol (10 mM), DiD (100 μM), and DONDI dye (40 μM) were prepared in chloroform/methanol (9:1, v/v). Giant unilamellar vesicles (GUVs) were formed in water containing 50 mM sucrose as previously described[1], using a Vesicle Prep Pro (VPP) instrument. GUV images were acquired on a Nikon A1R confocal microscope equipped with a 20× objective. DONDI probes were excited at 405 nm, and emission was collected using a GFP filter (500–550 nm). DiD was excited at 640 nm, and emission was collected between 663 and 738 nm. Individual vesicles were cropped from raw images, and vesicles exhibiting clear phase separation in the DiD channel were selected for analysis. Image analysis was performed using Fiji (ImageJ).[2]

**Molecular Dynamics Simulations and Analysis**

Three-dimensional structures of DONDI probes were generated using the CHARMM-GUI input generator, Ligand Reader & Modeler, and Membrane Builder modules.[3–5] CHARMM-compatible topology and parameter files were produced using the CHARMM36m force field for lipids and CGenFF-derived parameters[6,7] for all DONDI ligands. Membrane systems consisted of a POPC bilayer containing one DONDI molecule together with OOOTG (TAG) and DOGL (1,2-DAG) positioned in the same leaflet. The probe was oriented such that the naphthalimide moiety initially resided near the membrane–water interface. Systems were solvated with explicit TIP3P water, neutralized with counterions, and adjusted to 0.15 M NaCl under periodic boundary conditions. All molecular dynamics (MD) simulations were performed using GROMACS (v2022.1) with GPU acceleration. [8] Energy minimization was conducted using the steepest descent algorithm until convergence ($F_{max} < 1000$ kJ·mol$^{-1}$·nm$^{-1}$), with hydrogen-containing bonds constrained via the LINCS algorithm.[9] Equilibration proceeded through six restrained phases (initial NVT followed by semi-isotropic NPT), with gradual release of positional restraints on solute, membrane, and solvent components. Production simulations were performed for 300 ns using a 2 fs time step. The Verlet cutoff scheme was applied with neighbor list updates every 20 steps. Short-range electrostatic and van der Waals interactions were treated with a 1.2 nm cutoff, with van der Waals forces smoothly switched between 1.0 and 1.2 nm. Long-range electrostatics were calculated using the Particle Mesh Ewald (PME) method. Temperature was maintained at 298.15 K using the velocity-rescale (v-rescale) thermostat ($\tau = 1.0$ ps) with separate coupling groups for solute, membrane, and solvent. Pressure was controlled at 1 bar using the C-rescale barostat with semi-isotropic coupling ($\tau = 5.0$ ps; compressibility = $4.5 \times 10^{-5}$ bar$^{-1}$). Center-of-mass motion was removed every 100 steps. Coordinates were saved in compressed format every 100 ps. Unless otherwise stated, only the final 250 ns of each trajectory were used for quantitative analyses to ensure equilibration of probe insertion depth, orientation, and lipid–probe interactions. Trajectory analyses were performed using GROMACS utilities, VMD[10], and in-house scripts. Bilayer thickness was calculated from the average positions of phosphorus atoms in opposing leaflets. Probe immersion depth was defined as the distance between selected DONDI oxygen atoms and the bilayer center, defined as the midpoint between leaflet phosphorus planes. Naphthalimide tilt angles were determined from the angle between a vector defined by central scaffold carbon atoms and the membrane normal. Hydrogen bonds between DONDI, 1,2-DAG, TAG, and water were quantified using the VMD H-bond plugin[11] (donor–acceptor distance ≤ 3.0 Å; angle cutoff 30°). Lipid order parameters and probe orientation distributions were computed over every frame of the final 250 ns. Representative structures were visualized using UCSF Chimera (ver. 1.16).[12]

**Cell Culture**

NIH-3T3 fibroblasts (ATCC, USA) were rapidly thawed in a 37 °C water bath and immediately transferred to Matrigel-coated T-25 flasks (Corning). Cells were maintained in DMEM supplemented with GlutaMAX and 10% fetal bovine serum (FBS) at 37 °C in a humidified incubator with 5% $CO_2$ and grown to confluence. Confluent cultures were washed with PBS and detached using 0.05% trypsin–EDTA. Following centrifugation, cell pellets were resuspended in fresh culture medium and replated as needed.

For fluorescence imaging, cells were seeded onto glass-bottom, quad-divided, Matrigel-coated 35 mm dishes at a density of $1.0 \times 10^6$ cells/cm$^2$ and cultured under identical conditions. Experiments were performed 2–5 days after plating.

For HPLC analysis cells were seeded into Matrigel-coated T-25 flasks and grown to 70–90% confluence prior to treatment. Cells were incubated with 10 µM DONDI-5 for 24 h, then harvested by scraping and separated from the culture medium by centrifugation (500 × g, 5 min, 4 °C). Supernatants (culture medium) were stored at -20 °C until analysis. Cell pellets were washed once with fresh medium, centrifuged under the same conditions, and stored at -20 °C until HPLC analysis.

**Fluorescence Confocal Imaging**

Fluorescence imaging was performed on a Nikon AX confocal microscope equipped with an NSPARC (Nikon Spatial Array Confocal) detector and mounted on an Eclipse Ti2 inverted platform. Laser excitation lines of 405, 488, and 640 nm were used. A primary dichroic mirror (405/488/561/640 nm) was used for beam splitting. Sequential channel acquisition was employed to minimize spectral crosstalk. LysoView 640 was acquired in the first pass using 638 nm excitation with emission collected between 666–732 nm. In a second software-driven pass, BODIPY 493/503 was excited at 488 nm and DONDI at 405 nm, with emission for both channels collected between 502–546 nm. DONDI and BODIPY 493 signals were acquired sequentially within this pass to prevent excitation overlap. Emission windows were selected to minimize bleed-through between channels. Cells were incubated with the indicated DONDI probe (1 µM final concentration) for 1 h, washed with dye-free medium for the specified time period. Prior to imaging, cells were loaded with LysoView 640 (0.5 µM) and BODIPY 493/503, washed with dye-free medium, and transferred to an Oko-Lab environmental chamber mounted on the microscope stage. The chamber and 60x objective were pre-equilibrated to 37 °C, and imaging was performed at 37 °C under 5% $CO_2$ and saturated humidity. For each condition, three randomly selected fields of view were acquired at 1024 x 1024 pixel resolution. Z-stacks were collected with a 0.3 µm step size. Image processing and quantitative analysis were performed using ImageJ/Fiji.[2]

**Particle Intensity Analysis**

Fluorescence intensity across Z-stacks was first evaluated, and a representative optical section exhibiting high signal intensity was selected for further analysis. A median ("despeckle") filter was applied to reduce high-frequency noise prior to segmentation. Regions of interest (ROIs) were generated using the Trainable Weka Segmentation plugin[13] in Fiji/ImageJ. The classifier was trained on a series of images spanning both low and high signal-to-noise conditions to improve robustness. Classified outputs were converted to binary masks, from which ROIs were delineated. Particle intensity analysis was performed in Fiji/ImageJ using the original (unprocessed) images, with size thresholds restricted to 0.05–10 µm$^2$. For each ROI, mean fluorescence intensity and area were quantified. Equivalent particle diameters were calculated assuming circular geometry. Subsequent analyses were restricted to particles with diameters between 0.1 and 2 µm. Data from all time points were compiled and analyzed using GraphPad Prism (v10). Statistical significance was determined using the Wilcoxon matched-pairs signed-rank test.

**Colocalization Analysis**

Pearson's correlation coefficients (PCCs) were calculated using the EzColocalization[14] plugin in Fiji/ImageJ. Colocalization analysis was performed on single optical sections selected from each Z-stack based on maximal green-channel intensity. Costes' automatic thresholding was applied to minimize user bias. PCC values were determined in triplicate for each DONDI analog. Data from all time points were compiled and analyzed using GraphPad Prism (v10). Statistical significance was assessed using the Wilcoxon matched-pairs signed-rank test.

**Visualization and Drawing Software**

Reaction schemes and 2D structural formulas were prepared using MarvinSketch (ChemAxon Ltd., version 24.1, https://www.chemaxon.com). Some plots were obtained using RStudio (version 2026.01).

**HPLC Analysis**

Analytical HPLC was performed on a Shimadzu system (Japan) equipped with an LC-40D pump, a DGU-405 5-channel degasser, an SPD-M40 photodiode-array (PDA) detector, and an RF-20A fluorescence detector. Separations were carried out on an Agilent Poroshell 120 EC-C8 column (4.6 × 100 mm, 2.7 µm). The mobile phase consisted of solvent A, water containing 0.1% (v/v) trifluoroacetic acid (TFA), and solvent B, acetonitrile containing 0.1% (v/v) TFA. The method was isocratic at 90% B for 15 min, at a flow rate of 0.75 mL min$^{-1}$. Fluorescence detection was performed with excitation at 400 nm and emission collected at 540 nm.

**TLC analysis**

A DMSO stock solution of DONDI-5 was added to 0.1 M HEPES buffer containing 6 mM sodium cholate. Porcine carboxylesterase (porcine pancreas CEase; Worthington Biochemical, Cat. number LS004104) was then introduced to achieve a final enzyme concentration of 2.3 μM. The reaction mixture was incubated at 37 °C for 30 min. Aliquots were subsequently applied to a silica gel TLC plate and developed in chloroform/methanol (10:1, v/v). Bands were visualized under 395 nm illumination.

A1

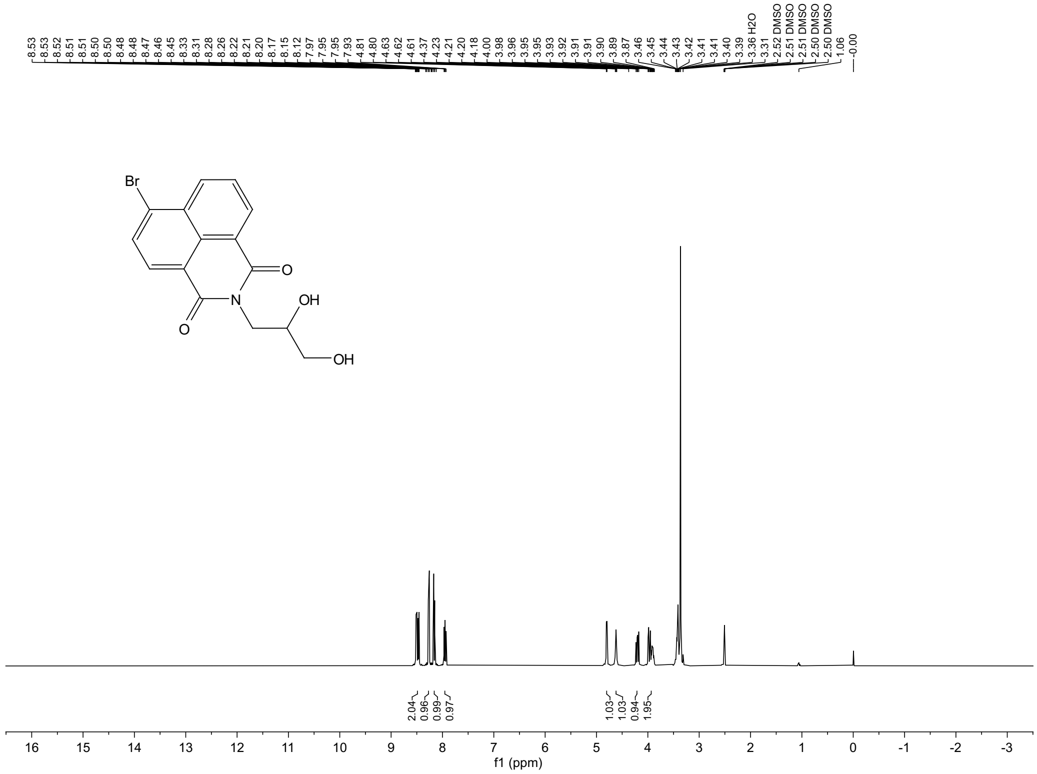
8.53
8.53
8.52
8.51
8.51
8.50
8.48
8.48
8.47
8.46
8.45
8.33
8.31
8.28
8.26
8.22
8.21
8.20
8.17
8.15
8.12
7.97
7.95
7.95
7.93
4.81
4.80
4.63
4.62
4.61
4.37
4.23
4.21
4.20
4.18
4.00
3.98
3.96
3.95
3.95
3.93
3.92
3.91
3.91
3.90
3.89
3.87
3.46
3.45
3.44
3.43
3.42
3.41
3.41
3.40
3.39
3.36 H2O
3.31
2.52 DMSO
2.51 DMSO
2.51 DMSO
2.50 DMSO
2.50 DMSO
1.06
-0.00
Br
O
N
OH
O
OH
2.04
0.96
0.99
0.97
1.03
1.03
0.94
1.95
16 15 14 13 12 11 10 9 8 7 6 5 4 3 2 1 0 -1 -2 -3
f1 (ppm)

A1

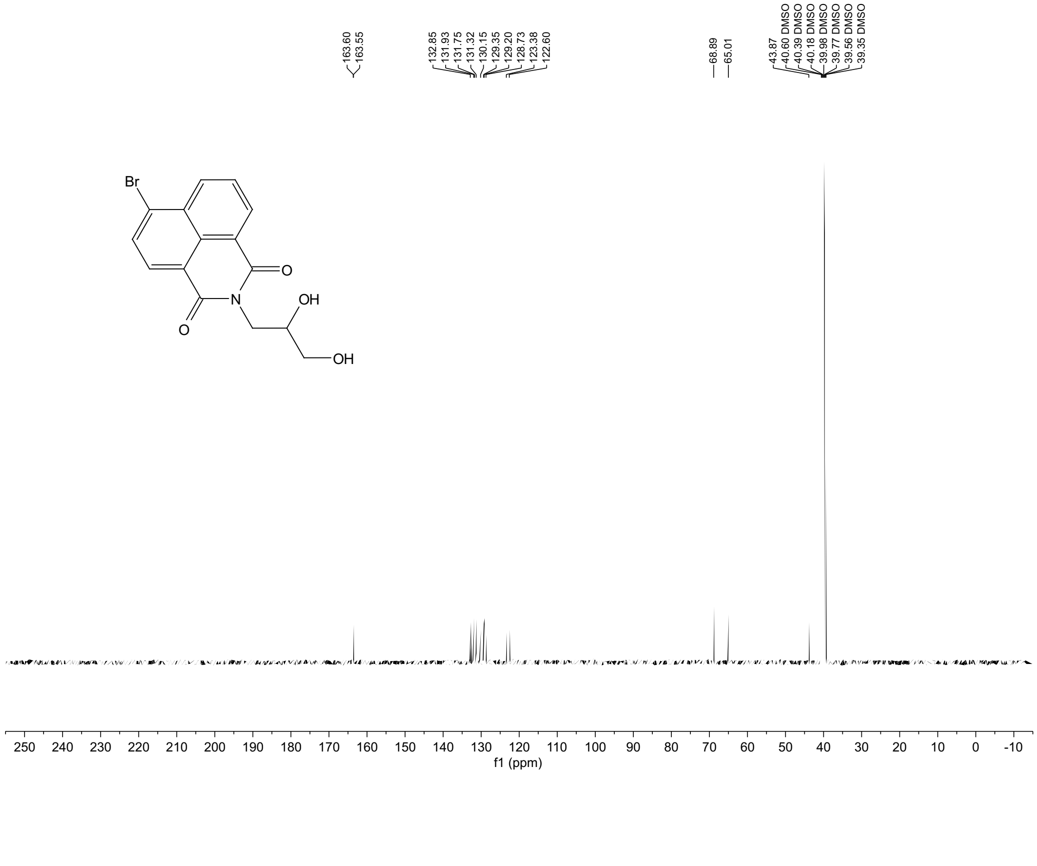
163.60
163.55
132.85
131.93
131.75
131.32
130.15
129.35
129.20
128.73
123.38
122.60
68.89
65.01
43.87
40.60 DMSO
40.39 DMSO
40.18 DMSO
39.98 DMSO
39.77 DMSO
39.56 DMSO
39.35 DMSO
Br
O
N
OH
O
OH
250 240 230 220 210 200 190 180 170 160 150 140 130 120 110 100 90 80 70 60 50 40 30 20 10 0 -10
f1 (ppm)

A2

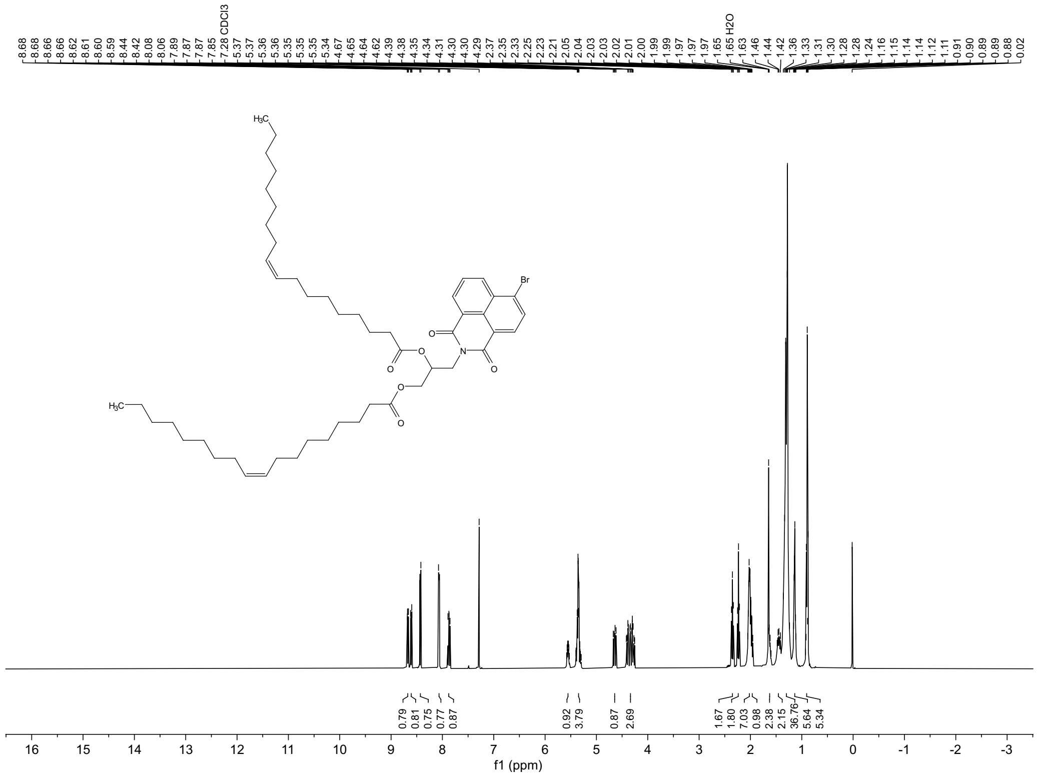
8.68
8.68
8.66
8.62
8.61
8.60
8.59
8.44
8.42
8.08
8.06
7.89
7.87
7.87
7.85
7.28 CDCl3
5.37
5.37
5.36
5.36
5.35
5.35
5.35
5.34
4.67
4.65
4.64
4.62
4.39
4.38
4.35
4.34
4.31
4.30
4.30
4.29
2.37
2.35
2.33
2.25
2.23
2.21
2.05
2.04
2.03
2.03
2.02
2.01
2.00
1.99
1.99
1.97
1.97
1.97
1.65
1.65 H2O
1.63
1.46
1.44
1.42
1.36
1.33
1.31
1.30
1.28
1.28
1.24
1.16
1.15
1.14
1.14
1.12
1.11
0.91
0.90
0.89
0.88
0.02
H3C
Br
O
N
0.79
0.81
0.75
0.77
0.87
0.92
3.79
0.87
2.69
1.67
1.80
7.03
0.98
2.38
2.15
36.76
5.64
5.34
16 15 14 13 12 11 10 9 8 7 6 5 4 3 2 1 0 -1 -2 -3
f1 (ppm)

A2

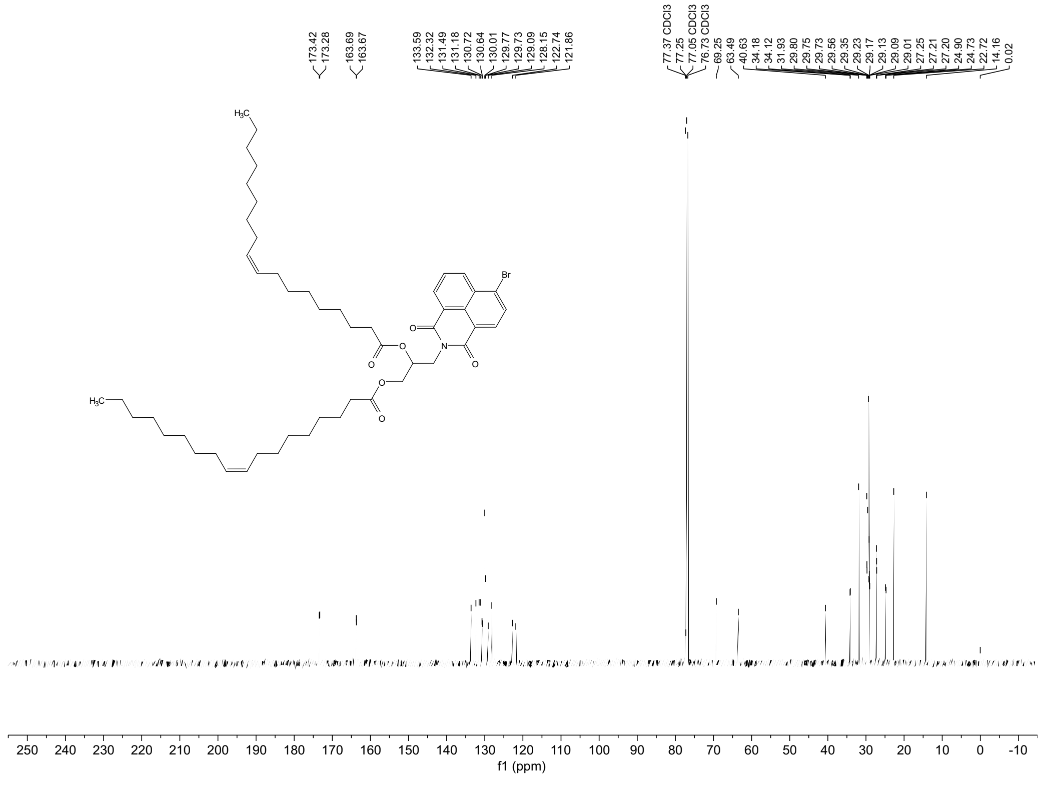
173.42
173.28
163.69
163.67
133.59
132.32
131.49
131.18
130.72
130.64
130.01
129.77
129.73
129.09
128.15
122.74
121.86
77.37 CDCl3
77.25
77.05 CDCl3
76.73 CDCl3
69.25
63.49
40.63
34.18
34.12
31.93
29.80
29.75
29.73
29.56
29.35
29.23
29.17
29.13
29.09
29.01
27.25
27.21
27.20
24.90
24.73
22.72
14.16
0.02
H3C
Br
O
N
250 240 230 220 210 200 190 180 170 160 150 140 130 120 110 100 90 80 70 60 50 40 30 20 10 0 -10
f1 (ppm)

DONDI-1

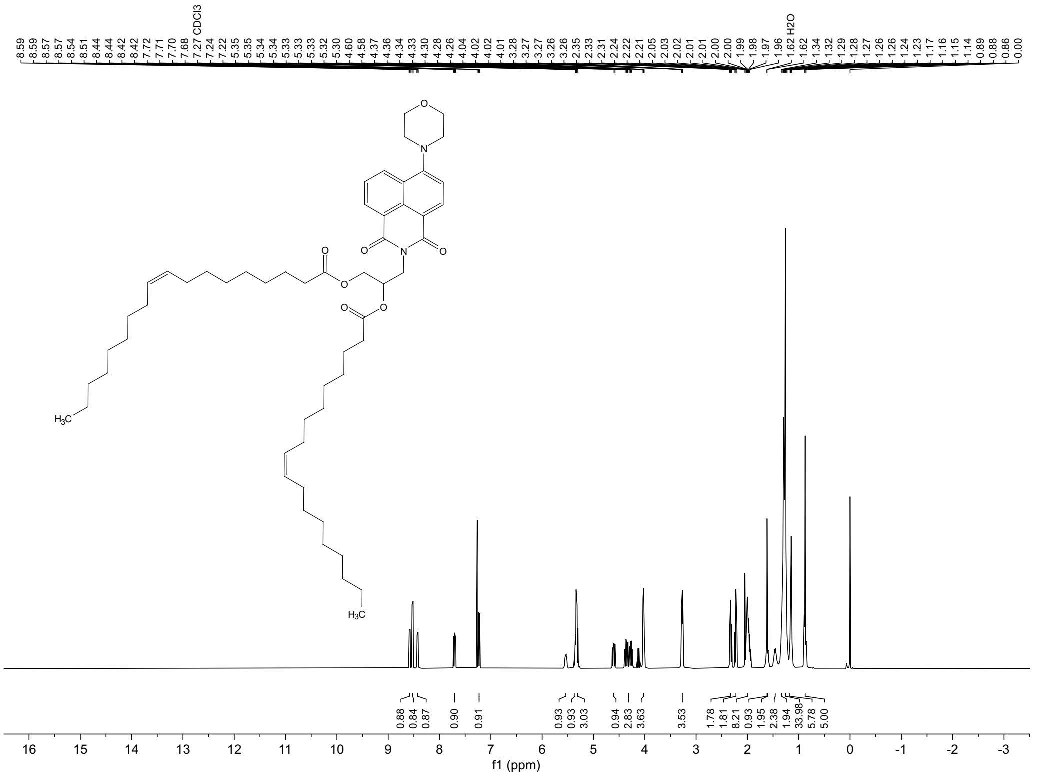

DONDI-1

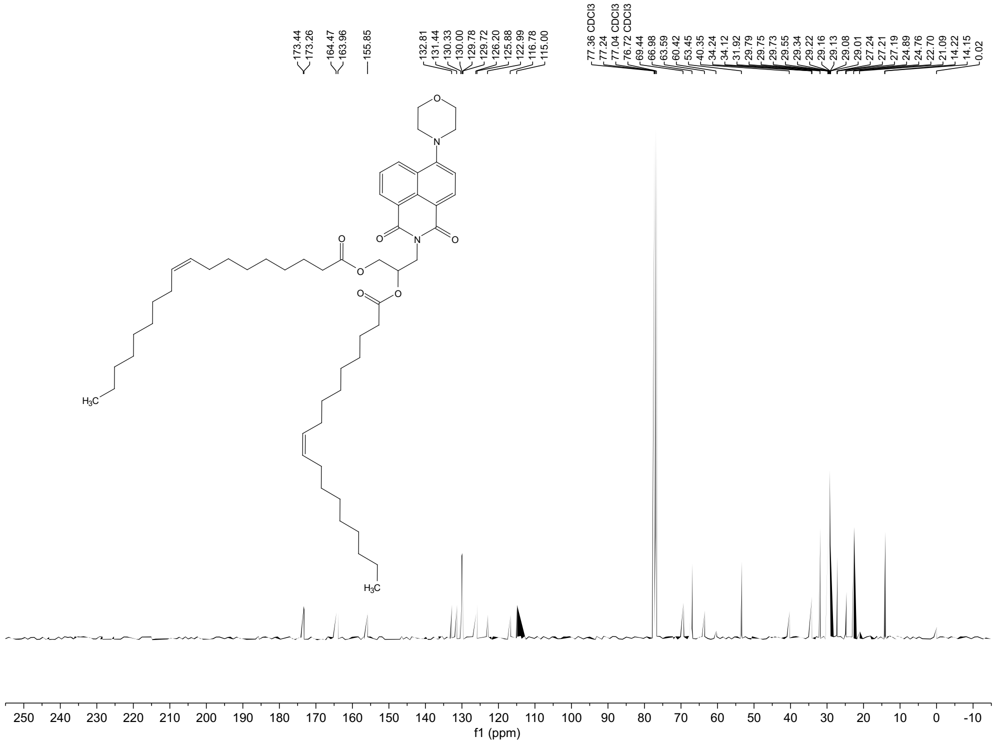

DONDI-3

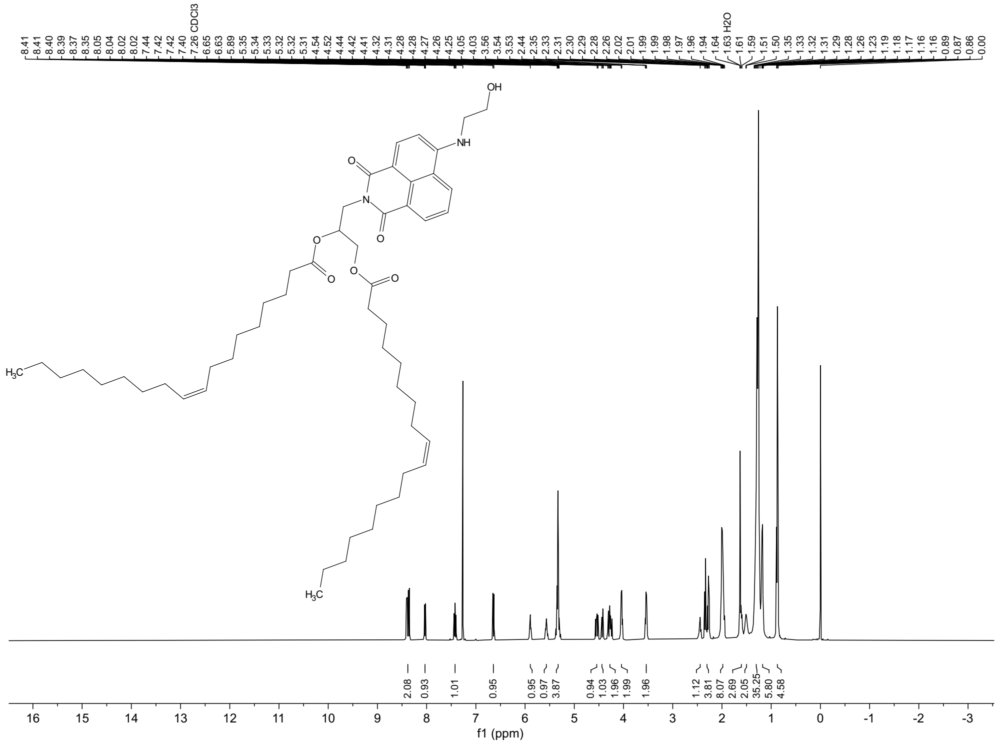

DONDI-3

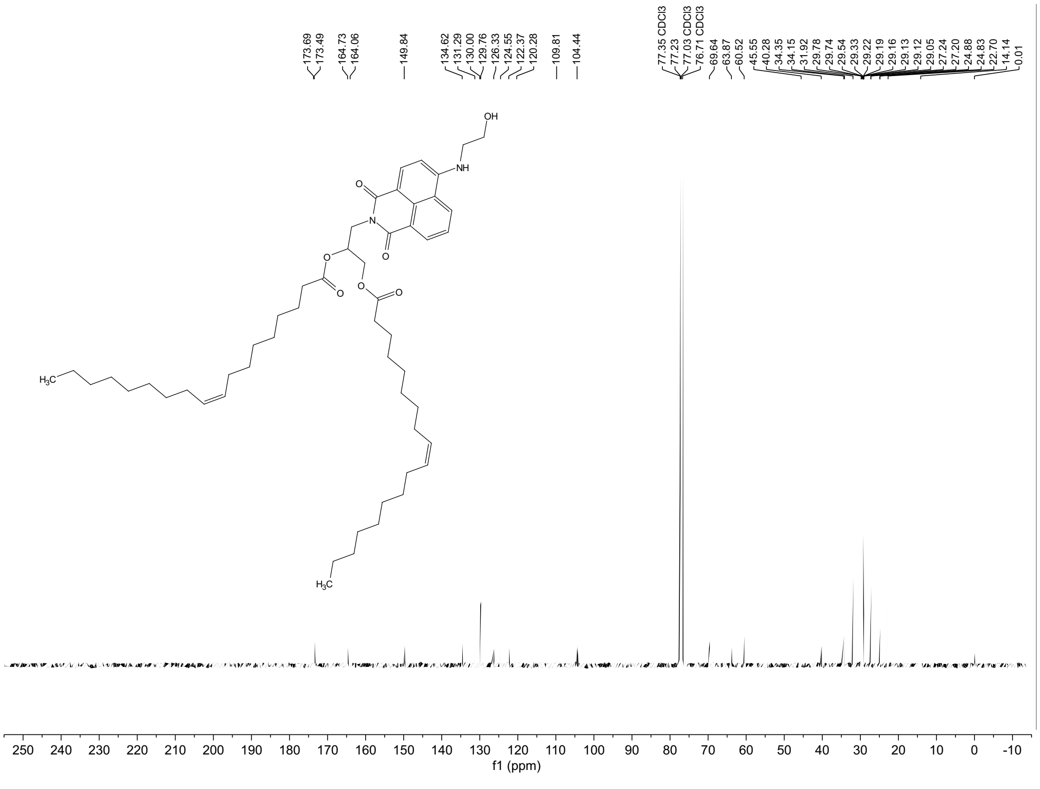

DONDI-4

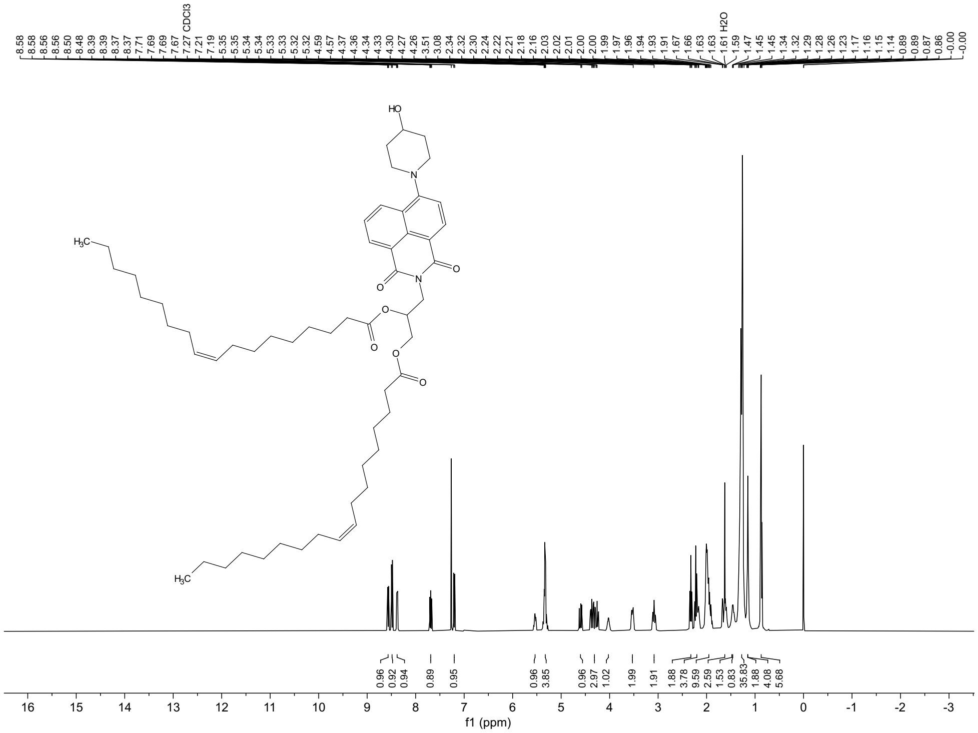

DONDI-4

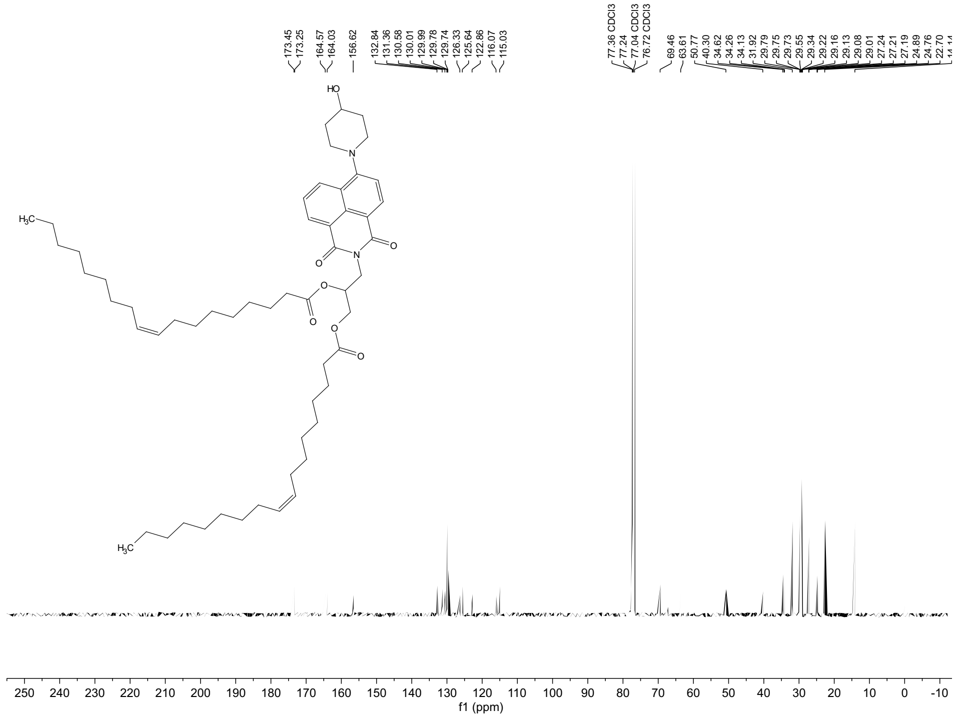

B1

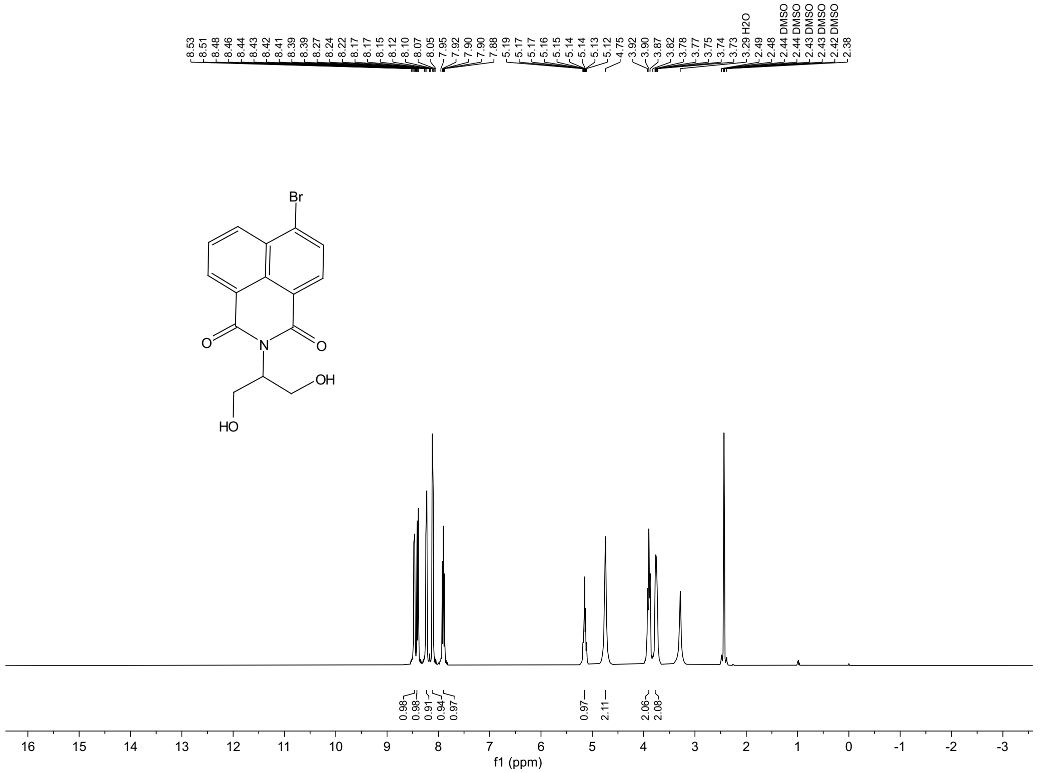
8.53 8.51 8.48 8.46 8.44 8.43 8.42 8.41 8.39 8.39 8.27 8.24 8.22 8.17 8.17 8.15 8.12 8.10 8.07 8.05 7.95 7.92 7.90 7.90 7.88 5.19 5.17 5.17 5.16 5.15 5.14 5.14 5.13 5.12 4.75 3.92 3.90 3.87 3.82 3.78 3.77 3.75 3.74 3.73 3.29 H2O 2.49 2.48 2.44 DMSO 2.44 DMSO 2.43 DMSO 2.43 DMSO 2.42 DMSO 2.38
Br
O
N
O
OH
HO
0.98 0.98 0.91 0.94 0.97 0.97 2.11 2.06 2.08
16 15 14 13 12 11 10 9 8 7 6 5 4 3 2 1 0 -1 -2 -3
f1 (ppm)

B1

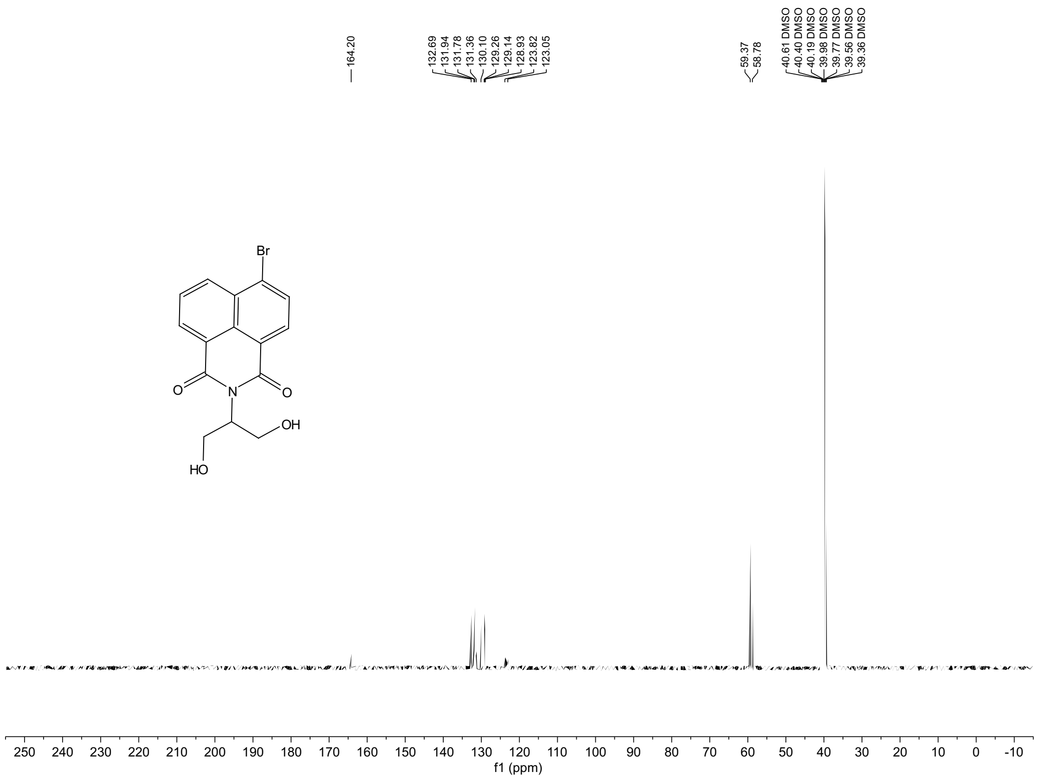
164.20
132.69 131.94 131.78 131.36 130.10 129.26 129.14 128.93 123.82 123.05
59.37 58.78
40.61 DMSO 40.40 DMSO 40.19 DMSO 39.98 DMSO 39.77 DMSO 39.56 DMSO 39.36 DMSO
Br
O
N
O
OH
HO
250 240 230 220 210 200 190 180 170 160 150 140 130 120 110 100 90 80 70 60 50 40 30 20 10 0 -10
f1 (ppm)

DONDI-5

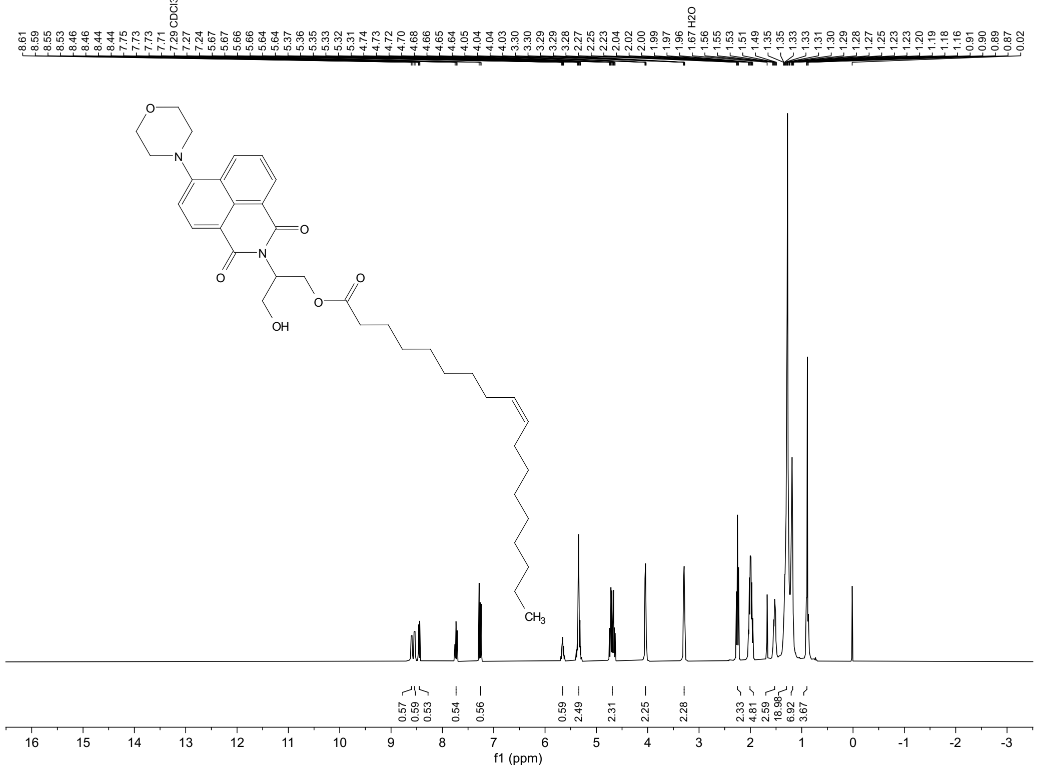

DONDI-5

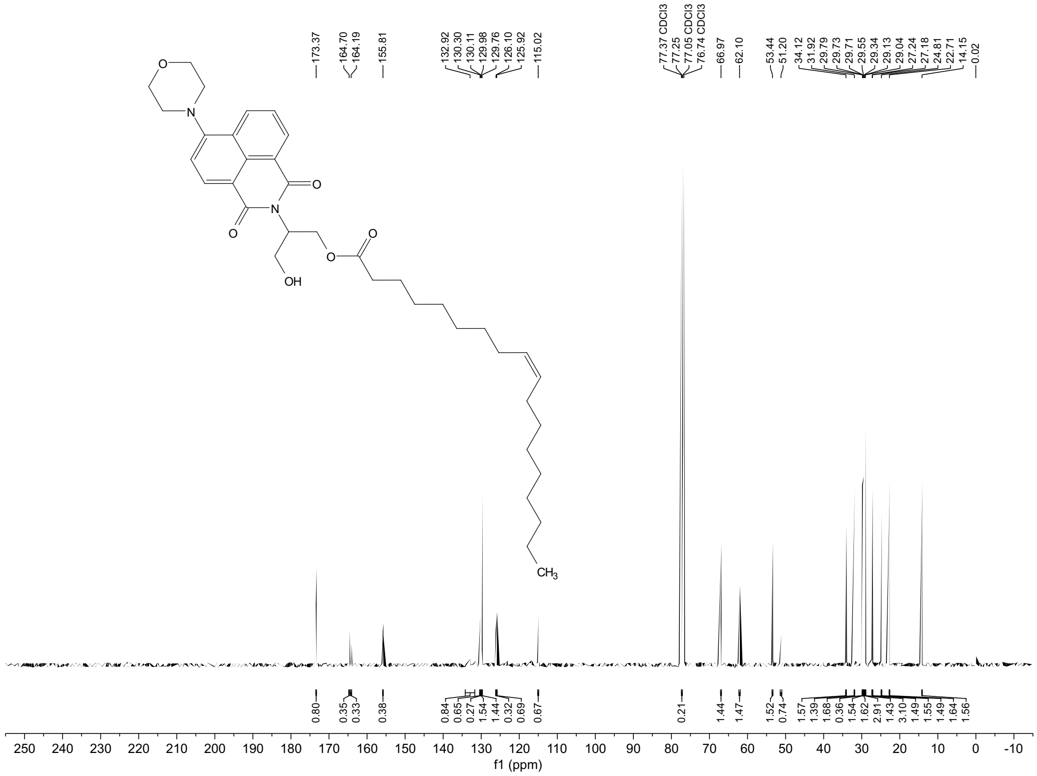

DONDI-1

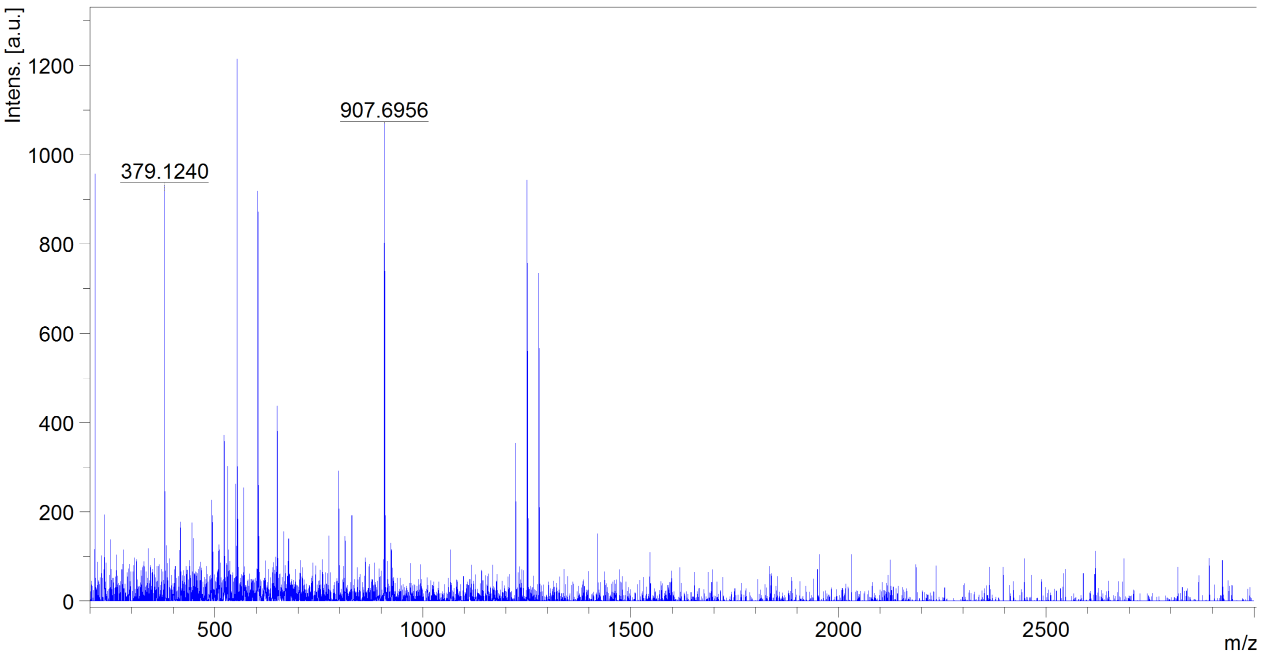

DONDI-3

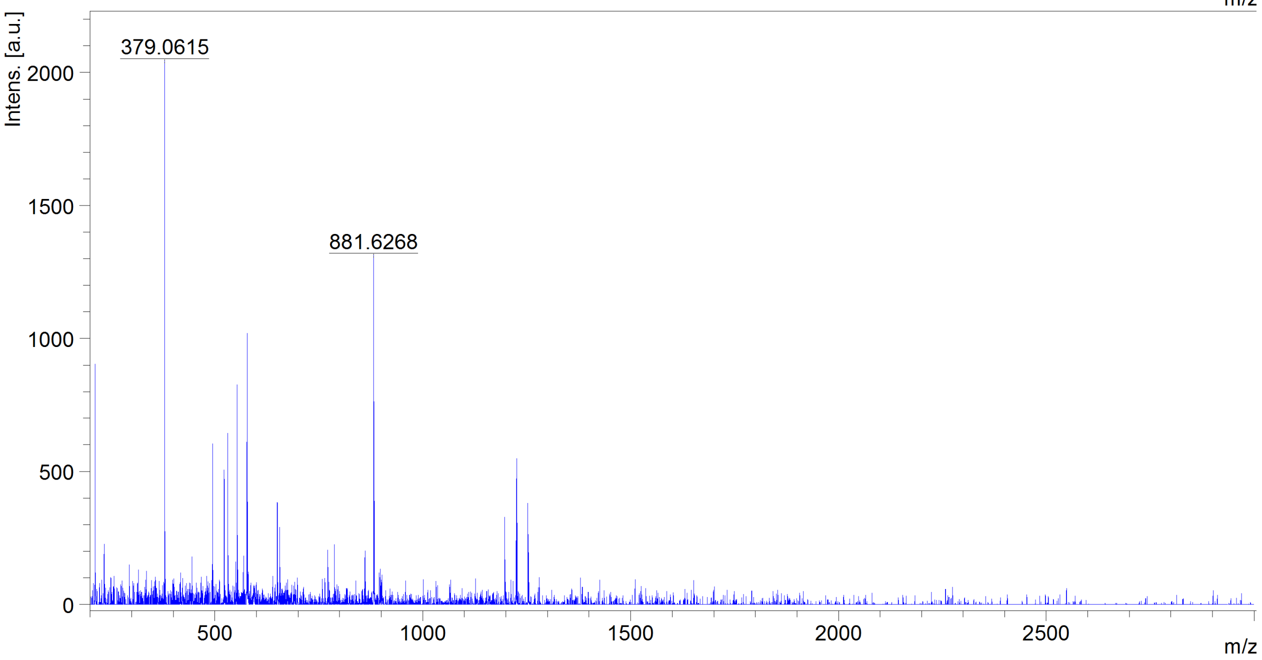

DONDI-4

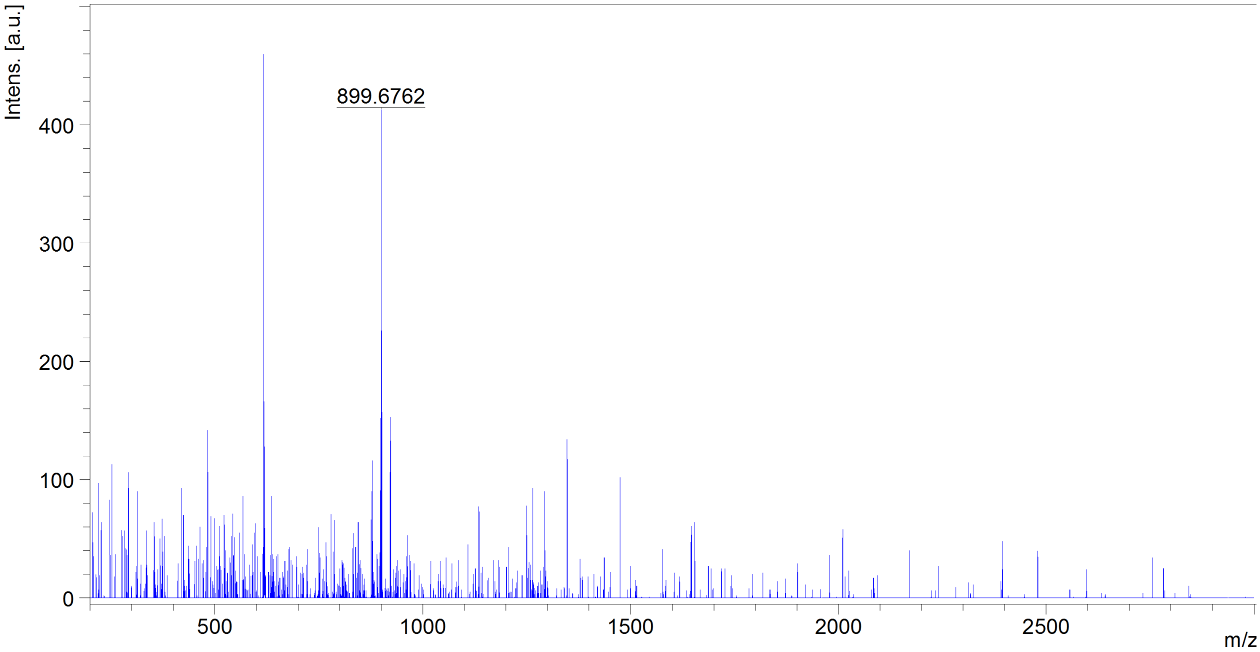

DONDI-5

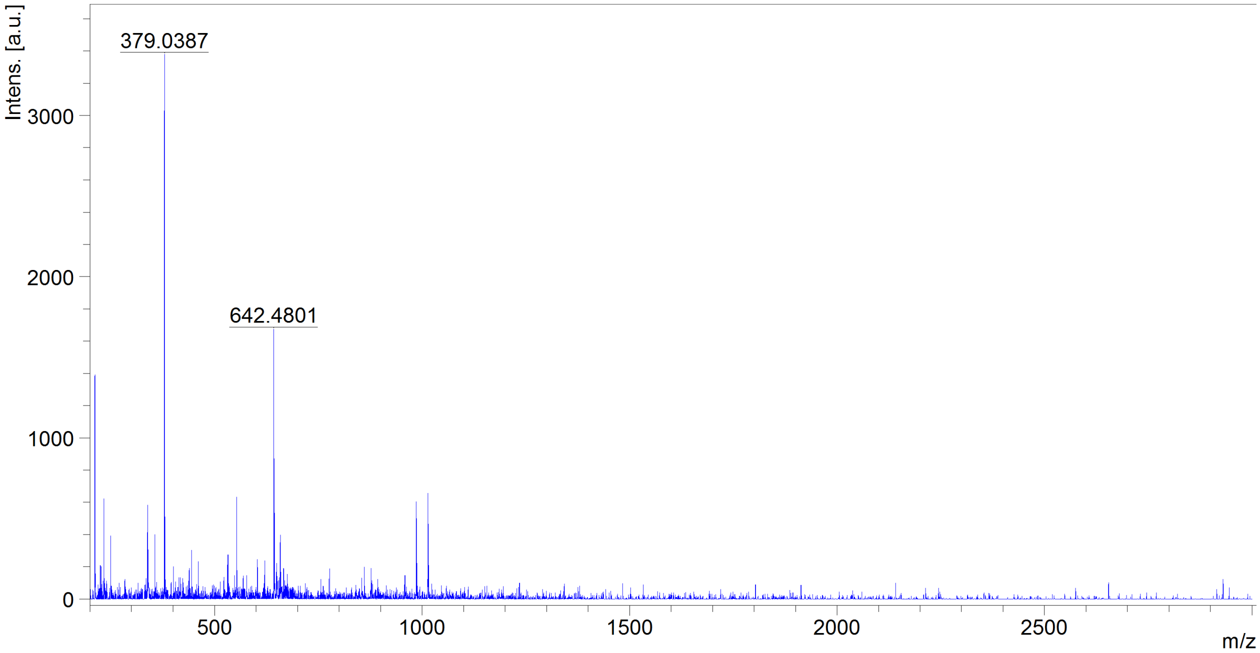
